# Engineering Synthetic Cell State Transitions by Harnessing Epigenetic Memory

**DOI:** 10.1101/2025.09.29.679324

**Authors:** Oscar A. Campos, Alessandro Migliara, Sarah R. Wasinger, Zhe Chen, Zoe L. Grant, Pilar Lopez, Benoit G. Bruneau, Satoshi Toda, Wendell A. Lim, Ricardo Almeida

**Author notes:** These authors contributed equally (Oscar A. Campos, Ricardo Almeida). Corresponding author (W.A.L.).

## Abstract

During development, cell-cell communication induces state transitions that are maintained by epigenetic regulation. Here, we harness endogenous epigenetic silencing machinery to engineer synthetic circuits that induce stable gene expression changes. Using synthetic Notch receptors that control the chromatin regulators KRAB and Dnmt3L, we developed input-sensing switches that induce self-sustained silencing of target loci. We used these modules to construct circuits in which combinatorial inputs direct a choice among multiple alternative states. These epigenetic switches can also be inverted to yield sustained activation of target genes. We demonstrate that these epigenetic switches can be used to drive morphological changes triggered by transient cell-cell interactions, but that remain stable over many cell divisions, as is observed in development. Additionally, these circuits can be used to drive the reprogramming of fibroblasts to a neuron-like state. These synthetic epigenetic circuits represent an important step towards engineering cell populations capable of coordinated multi-cell state decisions.

## INTRODUCTION

A cornerstone of multicellular life is the ability of progenitor cells to stably differentiate into specialized cell types. During development, cells undergo a complex series of state and fate transitions, which are induced and coordinated across the cell population by specific cell-to-cell signals.^1–3^ Such cell state transitions represent a committed change in gene expression that is mostly irreversible, barring unnatural or improbable events like reprogramming. Many of the signals that trigger state transitions are transient, happening at a particular stage of development,^4,5^ yet the induced cell state changes remain locked in, long after those developmental induction signals are gone. Thus, such committed changes allow cell fate trajectories to be shaped by the long-term history of inputs and interactions, including sequences of interactions that are well-separated in space and time. As cell engineering advances toward bottom-up construction of tissues and organs, it is becoming increasingly important to be able to direct similar types of committed cell state and fate transitions.^6–8^ Specifically, the ability to generate multiple alternative stable cell states from a common progenitor cell, and to do so in a way in which such states are irreversible, would be critical to program more complex multi-cellular structures and functions.^6,9^ Thus, a foundational goal for engineering cooperative multicellular systems is to develop synthetic gene circuits that can drive stable cell state transitions triggered by transient events in the cell’s history (Figure 1).^10–14^

**Figure 1.**
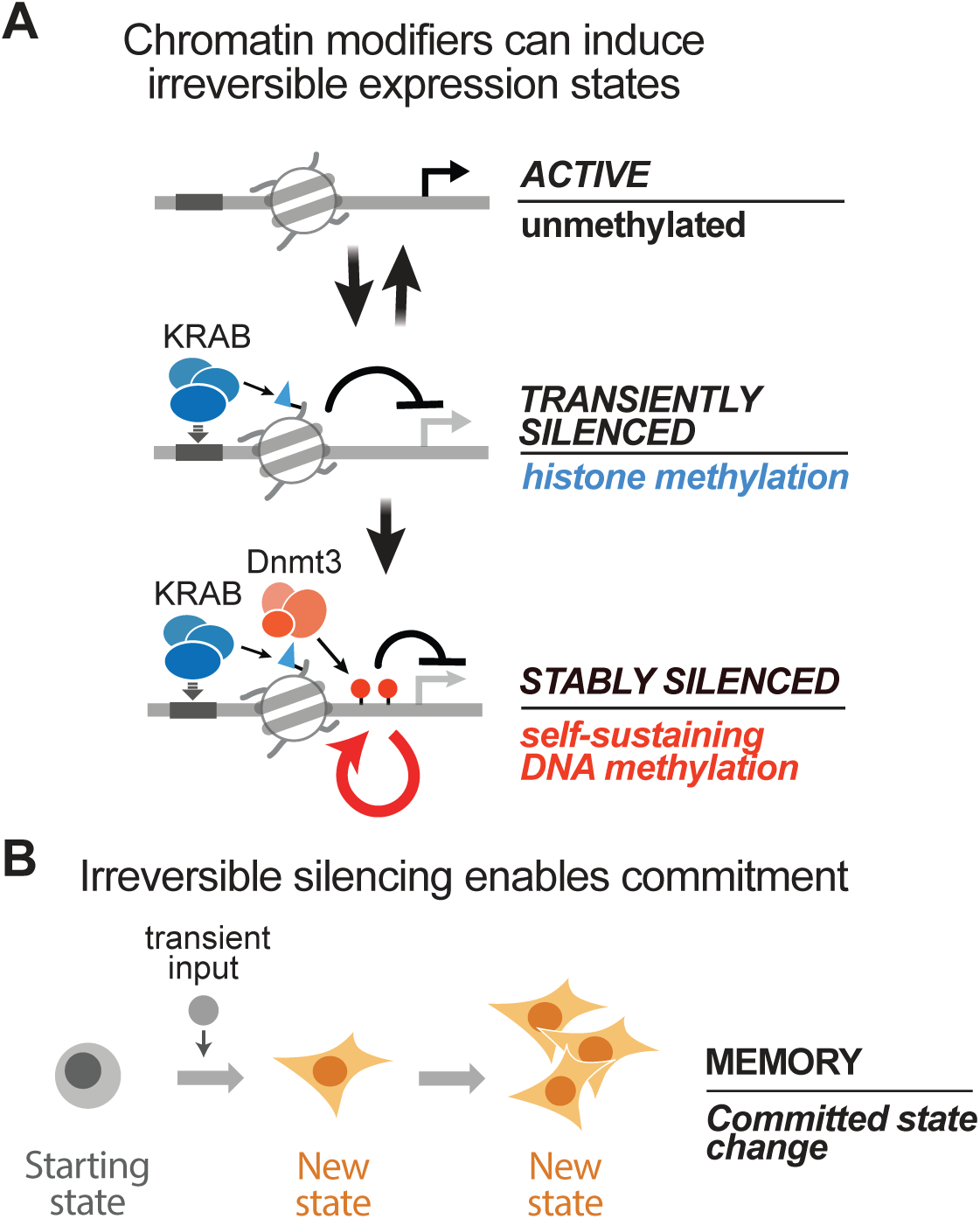
Epigenetic silencing by KRAB and Dnmt3 as a foundation for cell state commitment. **(A)** A three-state model describing a stepwise cooperation between KRAB domains and Dnmt3 to drive stable transcriptional silencing. KRAB domains, and the multimeric chromatin complexes they co-recruit, can establish a repressive heterochromatin domain to silence transcription in a reversible manner. In combination with Dnmt3, which establishes DNA methylation, the silenced state is locked in as repressive DNA methylation is effectively sustained over time and cell divisions. **(B)** A schematic representing an input signal that permanently switches cells to a new stable state regardless of whether the input persists. This scenario corresponds to the ability of Dnmt3 to establish epigenetic memory.

There is a long history in synthetic biology of engineering both prokaryotic and eukaryotic cells capable of adopting alternative stable states by constructing multi-transcription factor networks that contain mutual inhibition and positive feedback motifs.^15–17^ These bistable or multi-stable multi-transcription factor networks, however, can be challenging to integrate into more complex biological systems that would link them to both functionally relevant upstream inputs and downstream outputs. They also often lack the precision to drive selective transitions or to yield specific state distributions.

In eukaryotes, native cell state and fate transitions are also controlled by another type of mechanism: chromatin-based epigenetic modifications that provide stable control of gene expression at targeted loci in the genome.^18,19^ Epigenetic silencing can establish transcriptionally repressed states that are inherited across cell divisions.^20–23^ Thus, we postulated that it may be possible to engineer more modular and flexible synthetic cell state control circuits by harnessing the natural machinery of epigenetic regulation. A potent form of natural epigenetic regulation involves the cooperative effects of both histone modification and DNA methylation.^24–28^ Nucleosomes bearing histone H3 lysine trimethylation (H3K9me3) marks are recognized by heterochromatin protein 1 (HP1) which drives heterochromatin establishment and transcriptional repression.^29^ Through a series of protein-protein interactions, heterochromatin also recruits and activates the DNA methyltransferase 3 (Dnmt3) complex, which catalyzes cytosine methylation and further transcriptional silencing.^30^ Notably, DNA methylation is self-sustaining and inheritable as the DNA methylation pattern is efficiently maintained through replication by the maintenance methyltransferase, Dnmt1.^31–33^

Ectopic heterochromatin formation can be induced by artificially recruiting Krüppel associated Box (KRAB) domain fusion proteins to a target locus.^34–38^ While KRAB does inhibit gene expression, the heterochromatin state is relatively unstable (unlike DNA methylation) and alone does not yield stably silenced gene expression after removal of the KRAB signal (Figure 1A).^34,39–41^ On the other hand, cytosine methylation can be artificially induced by targeting the Dnmt3 complex, consisting of Dnmt3A, 3B, and 3L subunits (Figure 1A).^42–44^ However, recruitment of this complex alone is usually inefficient at silencing active promoters.^24,40^ Interestingly, simultaneous recruitment of KRAB and the Dnmt3 complex can generate silencing that is both efficient and stable across replication (Figure 1A). This synergy has been harnessed to stably silence target genes using a dCas9 fusion of KRAB, Dnmt3A and Dnmt3L.^24–26,28,45,46^ The precise dynamics by which KRAB-induced heterochromatin and Dnmt3-induced DNA methylation cooperate to yield efficient and stable silencing remains poorly understood.

Nonetheless, we realized that this potent self-sustaining silencing induced by KRAB and Dnmt3 might provide a powerful and more modular platform for engineering highly controllable circuits for cell state commitment. Such modularity could enable more complex programs that emulate natural developmental processes, including bifurcations into alternative states that are controlled by specific transient external stimuli, as well as sequential transition cascades (lineages) controlled by a sequence of transient stimuli.

Here, we harness the natural epigenetic machinery to engineer modular synthetic circuits that can: 1) establish commitment to stable cell state transitions, and 2) do so in ways that are induced by specific external cell-to-cell signals. To control both heterochromatin-induction and DNA methylation in response to external stimuli, we have linked KRAB and Dnmt3L domains to synthetic Notch (synNotch) receptors that require binding to cognate extracellular ligands to induce proteolytic release of the fused chromatin-regulating factors.^47^ We demonstrate that stable and efficient silencing of target genes requires co-incident stimulation of both the KRAB and Dnmt3 linked synNotch receptors. By linking these types of regulatory modules together, we can generate diverse memory behaviors including stable signal induced *activation* of target genes (in addition to *silencing*), and establishment of multi-state networks in which alternative paths can be selected via combinatorial input patterns. We demonstrate the ability to deploy such state switching modules to create self-organizing assemblies in which transient cell-cell interactions induce sustained morphological phenotypes, akin to what is observed in development. We also show that synthetic epigenetic memory circuits can be used to reprogram cells toward a different lineage, specifically the conversion of fibroblasts into a neuron-like state.

## RESULTS

### Inducible long-term epigenetic memory through synNotch triggering of KRAB and Dnmt3L activity

Synthetic circuits that can control stable cell transitions should have the following features: at least one output transcriptional unit than can exist in stable discrete “on” and “off” states, and a set of regulatory modules that can induce transitions between states in a directable manner. An ideal system can also enable cells to decide to undergo a transient or stable state change based on input identity.

To construct such a state transition circuit, we linked key chromatin regulators to synNotch receptors. Synthetic Notch receptors enable extracellular ligand binding to trigger proteolytic release of the intracellular effector domain, allowing it to enter the nucleus. In this case, as intracellular effector domains, we used DNA-targeted chromatin regulators, such that upon activation, they would bind to a designated target promoter. We first constructed a synNotch receptor with an extracellular anti-CD19 single-chain variable fragment^48^ (with a Myc epitope tag) linked to an intracellular TetR DNA binding domain fused to the Kox1 KRAB domain.^38,49^ This myc-anti-CD19-Notch-TetR-KRAB receptor was designed, in principle, to induce KRAB mediated silencing at a target locus, upon stimulation with the extracellular ligand CD19. We constructed a second synNotch receptor with an extracellular anti-GFP nanobody^50^ LaG16 (with an HA epitope tag) linked to an intracellular Gal4 DNA binding domain fused to the regulatory subunit of the mouse Dnmt3 complex, Dnmt3L (D3L) (Figure 2A).^51–53^ The resulting HA-LaG16-Notch-Gal4-D3L receptor was designed, in principle, to recruit the Dnmt3 DNA methylation complex to the target locus, upon stimulation with the extracellular ligand, GFP (Figures 2A and S1A). In preliminary studies, we found that recruitment of the D3L subunit alone was the most compact and sufficiently effective approach to induce targeted epigenetic memory. Our finding is in agreement with prior studies that artificially leverage D3L to recruit endogenous Dnmt3A/B,^28^ including a recent advance of the CRISPR-based epigenetic silencing system, CRISPRcharm, that devised a compact silencer construct containing an engineered histone H3 tail and truncated D3L domain.^28,54^ Altogether, these two receptors should enable independent sensing of two different inputs that are then coupled to discrete DNA targeting by the two different classes of chromatin regulators (Figure 2A).

**Figure 2.**
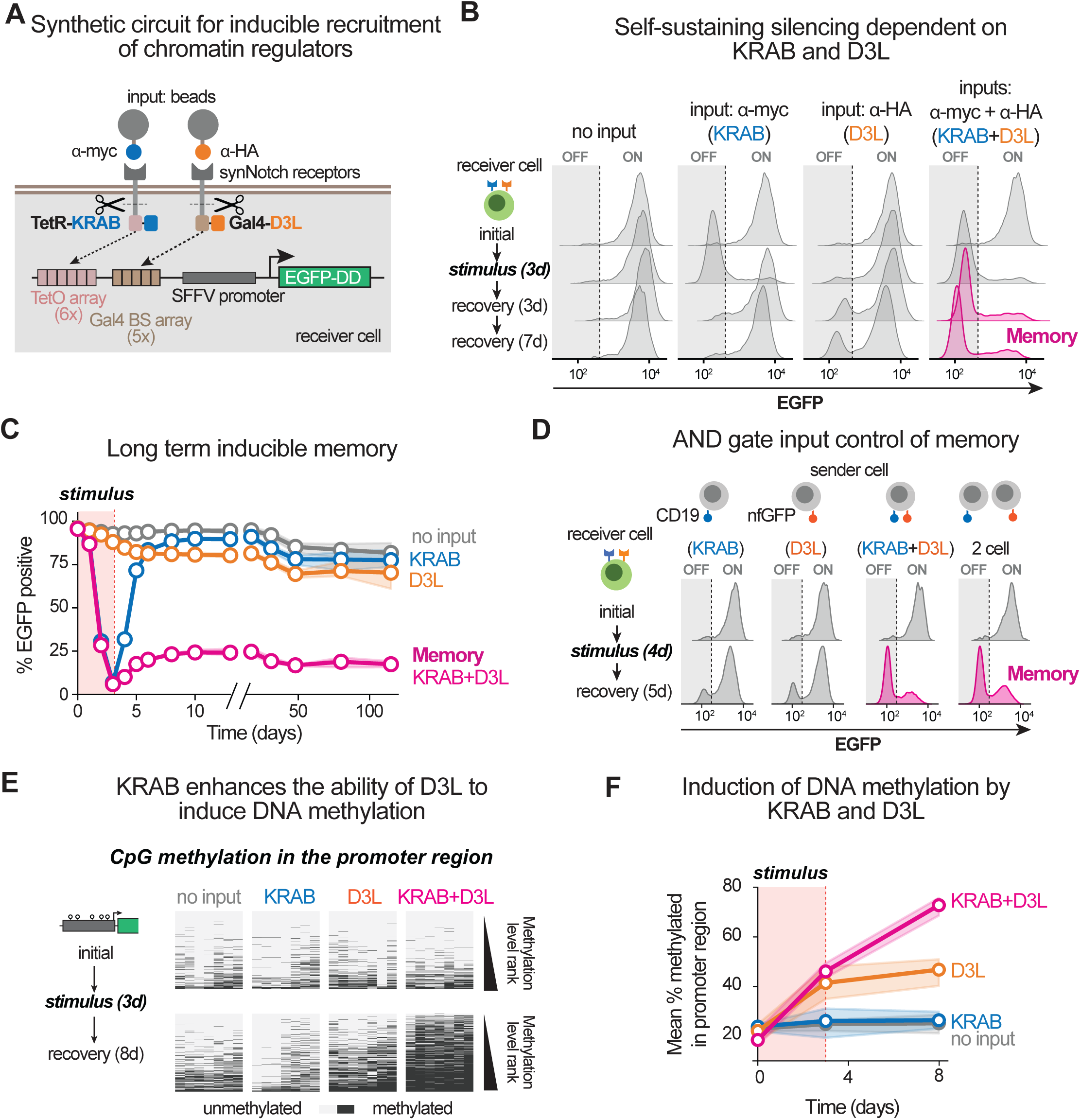
Inducible epigenetic memory through synNotch control of KRAB and Dnmt3L regulation. **(A)** Design of a receiver cell engineered to contain a genomically integrated synthetic circuit that enables the inducible site-specific recruitment of KRAB and D3L domains to a target locus. **(B)** Flow cytometry distributions of receiver cells incubated with magnetic beads as indicated. Cells were stimulated for 3 days and allowed to recover for up to 7 days by removal of beads and in the presence of DAPT. Distributions are representative of 3 technical replicates. **(C)** Time course of an experiment similar to (B), but in which cells were followed for 112 days after a stimulus pulse. The graph shows the percent of cells expressing EGFP, as determined by the threshold indicated by the dotted lines in (B). Data are from 3 technical replicates. Confidence intervals (95%) are the shaded areas. **(D)** Same as (B) except for cells co-cultured with ligand-presenting sender cells as indicated. Cells were co-cultured for 4 days to trigger synNotch receptors (distributions now shown) and allowed to recover for 5 days. **(E)** Methylation status of 7 CpG sites near the 3’ end of the engineered SFFV promoter. Rows are individual sequenced PCR amplicons ranked by total methylation level. Cells were stimulated for 3 days (plots not shown) and allowed to recover for up to 7 days by removal of beads and in the presence of DAPT. Plots are representative of 3 technical replicates. **(F)** Time course of average CpG methylation for 20 cytosines across the engineered SFFV promoter. Cells were transiently incubated with magnetic beads as in (B) and recovered in the presence of DAPT. Confidence intervals are the shaded areas.

Correspondingly, we constructed target promoters whose transcriptional state could be controlled by the two synNotch-gated chromatin regulators (Figure 2A). We placed binding site arrays for the TetR and Gal4 DNA binders upstream of the strong constitutively active spleen focus-forming virus (SFFV) promoter (Figure 2A). Altogether, this synNotch-promoter system constitutes a switching module that could control the expression of any payload gene. We used enhanced green fluorescent protein (EGFP) as a reporter, whose expression would reflect the chromatin environment and transcriptional state of the promoter (Figure 2A). Lastly, we fused the FKBP-DD degron^55^ (DD) to the EGFP reporter protein to minimize the delay between transcriptional repression and fluorescence decay (Figure 2A).

We generated a “receiver” cell using the mouse L929 fibroblast cell line, introducing the synNotch receptor and target promoter circuits via lentiviral integration. We also generated a corresponding set of “sender” L929 cells that express cell surface CD19 and/or non-fluorescent GFP mutant (nfGFP) ligands,^56^ which could trigger one or both of the synNotch receptors. The synNotch receptors could also be triggered by stimulating with either anti-Myc or anti-HA beads that recognize the extracellular epitope tags.

Receiver cell stimulation by incubation with anti-Myc beads (Figure 2A and S1B) or by co-culture with CD19-presenting sender cells (Figure S1C) should induce KRAB domain recruitment to the target reporter promoter. Indeed, we observed that EGFP expression was substantially silenced within 3 days of stimulation, with about 80-90% of cells in the polyclonal population switching to an EGFP^low^ state (Figure 2B, 2C, and S1C). However, this silencing is transient as EGFP expression is largely recovered following inhibition of synNotch by addition of the gamma-secretase inhibitor, DAPT^57^ (Figure 2B, 2C, and S1C). This is consistent with previous findings that KRAB-mediated transcriptional silencing is largely reversible and is consistent with substantial reversal of heterochromatin by activating factors recruited to the SFFV promoter.^34,39–41^ In contrast, stimulation with anti-HA beads (Figure 2A and S1B) or co-culture with nfGFP sender cells (Figure S1C), which should recruit the Dnmt3 complex to the reporter promoter, resulted in a much lower percentage of cells silencing EGFP (Figure 2B, 2C, and S1C). This is consistent with previous findings that Dnmt3-mediated transcriptional silencing is, by itself, ineffective at silencing strong, constitutively active promoters.^40^

Crucially, we observed that concurrent stimulation of synNotch-KRAB and synNotch-D3L receptors results in the same potent EGFP silencing as KRAB alone but, importantly, additionally results in commitment to the EGFP^low^ state in almost all cells (Figure 2B, 2C, and S1C). Inhibition of the synNotch stimulus (by DAPT) does not restore EGFP expression, indicating a switch to a self-sustaining silenced chromatin state and epigenetic memory. The target gene remained stably silenced for longer than 3 months or about 100 cell generations (the longest timepoint we measured). Notably, memory could be achieved via concurrent stimulation with any input combination that stimulated both KRAB and D3L synNotch receptors, including anti-Myc and anti-HA beads (Figure 2B and 2C), co-culture with a sender cell that co-expresses both CD19 and nfGFP ligands (Figure 2D and S1C), or with simultaneous co-culture with a combination of two different sender cell types, each expressing one of the ligands (Figure 2D). Thus, the dual receptor system not only drives effective silencing and memory but also maintains the flexibility of generating transient vs stable switching in a single cell type depending on the input combination sensed.

Altogether, these results are consistent with a model in which Dnmt3-mediated DNA methylation is greatly enhanced by KRAB mediated silencing. In other words, heterochromatin induction is required to effectively initiate silencing, but DNA methylation makes silencing stable (i.e., committed). To corroborate this general hypothesis, we examined the changes in both DNA methylation (Figure 2E and 2F) and heterochromatin (Figure S1D) at the promoter region. Specifically, we examined the dynamics of CpG methylation (via cytosine conversion and sequencing) as well as H3K9me3 modification, a canonical mark for heterochromatin (via ChIP-qPCR). These modifications were tracked following transient synNotch stimulation (3 days) and a recovery phase (5-10 days). Stimulation of synNotch-KRAB alone induced an increase in H3K9me3 methylation, but this increase was transient (Figure S1D). Notably, synNotch-KRAB activation alone does not lead to any increase in CpG methylation. However, synNotch-KRAB activation,when combined with synNotch-D3L activation, leads to a synergistic increase in promoter CpG methylation (Figure 2E and 2F). Crucially this DNA methylation remained elevated even after input removal. Altogether these findings are consistent with a model where the ability of KRAB to induce heterochromatin (and silencing) strongly primes Dnmt3’s activity of inducing CpG methylation. Subsequently, elevated CpG methylation can sustain the silenced state even if H3K9me3 is lost (Figure S1D). Neither Dnmt3 nor CpG methylation stabilizes H3K9me3. Moreover, these results indicate that DNA methylation and not H3K9me3 is the crucial molecular determinant of self-sustaining memory.

We further tested this notion by treating the population of cells that had received dual synNotch-KRAB and synNotch-D3L stimulation with either the DNA methyltransferase inhibitor 5-aza-2’-deoxycytidine (5-aza)^58^ or the TET1 methylcytosine dioxygenase enzyme, which is a major factor in DNA demethylation.^59^ We observed reactivation of EGFP in about 75% of cells in both cases (Figure S1E and S1F), supporting the critical contribution of DNA methylation to the epigenetic memory observed in this system. Interestingly, cells in which EGFP was reactivated by TET1 expression can be re-silenced once more by another synNotch input pulse, suggesting a successful memory reset and re-establishment of the initial chromatin state. This is in line with previous epigenetic editing approaches.^24^ In summary, we show that synNotch receptors can be used to control epigenetic regulators, allowing epigenetic memory to be induced by external cell-to-cell signals. These synNotch circuits constitute minimal switching modules with which to engineer more complex cell state transitions.

### Transcription state bifurcation emerges from synthetic memory switch

Memory switches are predicted to yield bistability, which can be observed as all-or-none bifurcation of individual cells into discrete ON or OFF states.^19,22,60,61^ Thus, we wanted to examine whether these epigenetic switches yielded bistability. Lentiviral transduction can lead to heterogeneity of expression outcomes within a bulk cell population (probably due to reporter integration into variable chromatin environments). Thus, we first sought to characterize a panel of isogenic single cell clones derived from the polyclonal population harboring TetR-KRAB synNotch, Gal4-D3L synNotch, and the EGFP response element (Figure 2A). In response to a 3-day synNotch stimulus pulse, ∼73% of clones transiently silenced EGFP expression with KRAB input alone, and ∼85% of these clones displayed evidence of epigenetic memory with KRAB and D3L input (classified as >1/3 of the population maintaining a stable silenced state). This broadly recapitulated the response of the polyclonal population, including also the relative lack of response to D3L input alone (Figure S2A). Interestingly, for many of the clones in this system in which there was fractional stable silencing, the population bifurcated into either fully active or fully silenced states, instead of showing a distribution of intermediate expression levels (Figure S2A and S2B).

We characterized in more detail three clonal lines whose circuit performance was broadly representative of the bulk population. We used these clonal cell lines to test whether the epigenetic circuit could produce population bifurcation in response to different input strengths. We set up a series of stimulation regimens where receiver cells were subjected to stepwise increases in dual stimulation duration, ranging from 0.5 days to 3 days (Figure S2C). We observed a non-linear time-dependence for generating stably silenced cells (Figure 3A, S2C, S2D, and S2E), wherein a stimulus duration increase from 1 to 1.5 days resulted in more than 2-fold increase in the proportion of silenced EGFP^low^ cells. With 2 days of stimulation, >90% of the population stably switched to the silenced state (Figure 3A, S2C, and S2D). Importantly, stimulus duration did not correlate with the final mean fluorescence intensity following the recovery period but instead resulted in obvious bifurcation between silenced EGFP^low^ and active EGFP^high^ cells (Figure 3A).

**Figure 3.**
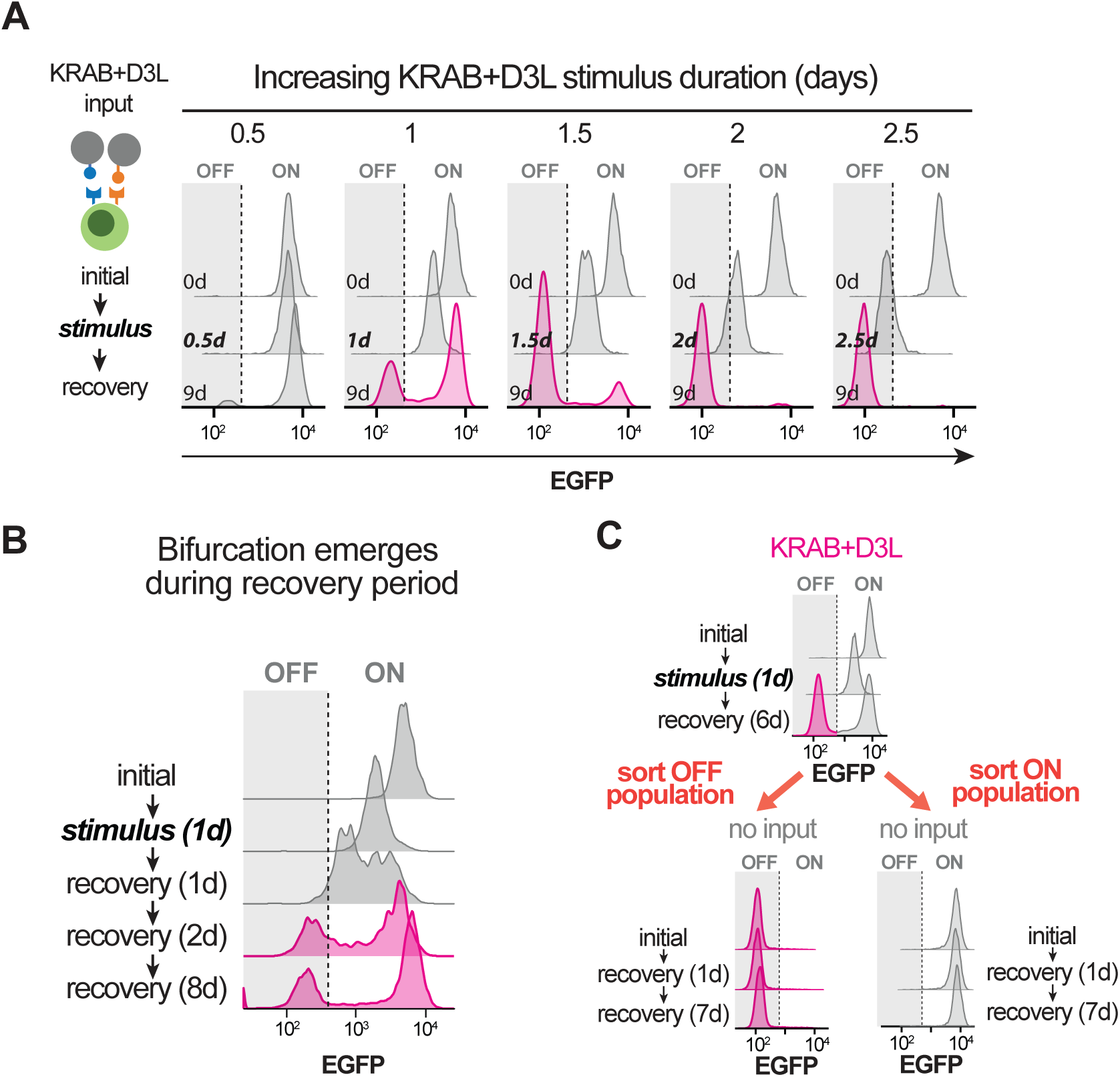
Synthetic memory switch results in bifurcation at single cell level. **(A)** Flow cytometry distributions of receiver cells stimulated for varying amounts of time, as indicated. Distributions are representative of 3 technical replicates of a single cell clone. **(B)** Same as (A), except for cells stimulated for only 1 day. **(C)** Same as (A), except that cells with high and low EGFP expression following KRAB+D3L stimulation were sorted by FACS at the end of a first recovery period. Cells were then followed for an additional week without input.

Intriguingly, short stimulus durations, such as the 1-day pulse, resulted in a silencing response characterized by two phases. Initially, the population uniformly represses EGFP (Figure 3B and S2F). However, over time and after the end of the stimulation period, the population resolves into distinct silenced and active subpopulations (Figure 3B and S2F), which is consistent with stochastic all-or-none state transitions observed in natural differentiation processes.^62–64^ Cells in which KRAB only was stimulated showed no bimodal population dynamics (Figure S2G), suggesting the dependence on D3L and epigenetic memory for this dynamic bifurcation behavior. Mechanistically, this also suggests that heterochromatin and DNA methylation undergo a slow autoregulatory spreading process.^65^

We further tested if population bifurcation is the result of cells committing to the silenced state as opposed to cells very rapidly switching between states following synNotch stimulation. We sorted EGFP^low^ and EGFP^high^ populations using FACS and followed them for an additional week. Crucially, cells maintained their expression state over time. Cells that were silenced did not recover the ON state, consistent with a state commitment (Figure 3C). Additionally, we find that EGFP^high^ cells did not spontaneously silence EGFP (Figure 2C), yet they bifurcated again following an additional short pulse, suggesting that the unsilenced cells maintain responsiveness to induced silencing despite prior stimulus encounters (Figure S2G).

### Co-incident KRAB and D3L stimulation is critical for effective state switching

These findings demonstrate the requirement for stimulation of both synNotch-KRAB and synNotch-D3L receptors to yield an epigenetic memory response, which is consistent with previous studies that have harnessed the interplay between heterochromatin and DNA methylation.^24,25^ However, the two-input circuits now provide an opportunity to address unanswered questions about the temporal relationship between KRAB and Dnmt3 silencing modules.

To gain insight into the temporal dependencies of the KRAB and D3L synNotch silencing system, we took advantage of the modularity of the circuit to test a series of stimulation regimens where we maintained a constant saturating stimulation duration of 4 days for both factors but varied the relative start time of each (Figure 4A, 4B, and S3A). Correspondingly, this set of experiments systematically alters the overlap time between KRAB and D3L stimulation and includes regimens where synNotch-D3L was stimulated entirely before or entirely after synNotch-KRAB. Overall, as the duration of temporal coincidence between KRAB and D3L stimulation is increased, ranging from no overlap to a full 4-day overlap, the proportion of cells that stably silenced EGFP also increased (Figure 4A, 4B, and S3B). Similar to the result obtained with short stimulus pulses, shorter temporal overlaps produced bifurcation of the cell population between OFF and ON states (Figure S3B). Notably, partial overlaps still resulted in a high percentage of the population stably silencing EGFP (Figure 4A, 4B, and S3B).

**Figure 4.**
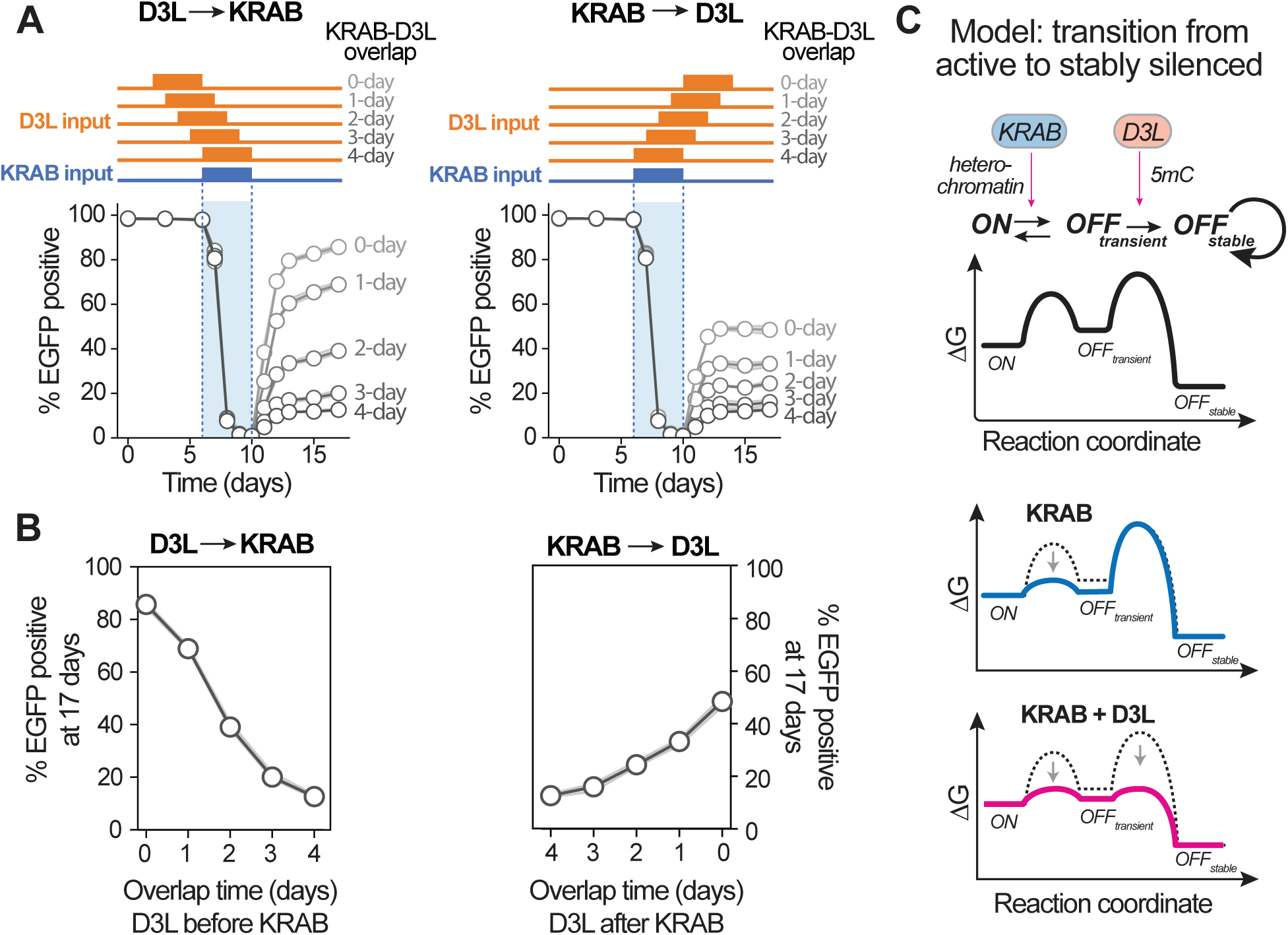
Coincidence and order of KRAB and D3L stimulation is critical for effective state switch. **(A)** Time courses of the percent of cells expressing EGFP for varying stimulation regimens. The blue shaded area indicates the time of KRAB stimulation, with the staggered time of D3L stimulation shown as orange bars above. The times of KRAB-D3L overlap are also indicated. Data for all curves are from 3 technical replicates of a single cell clone. Confidence intervals are mostly too small to be visible. **(B)** Summary of the experiment in (A) showing the percent of cells that stably silenced EGFP as a function of the overlap time between KRAB and D3L stimulation. The left side represents regimens where D3L stimulus preceded KRAB, and the right side is where KRAB stimulus preceded D3L. Data are from 3 technical replicates of a single cell clone. Confidence intervals are mostly too small to be visible. **(C)** Model describing the stepwise transition of a promoter from transcriptionally active (*ON*) to stably silenced (*OFF_stable_*). 5mC: 5-methylcytosine.

Nonetheless, the order of KRAB vs D3L stimulation was also crucially important. Stimulation regimens in which KRAB input preceded D3L resulted in greater proportion of stable silencing than the inverse, where D3L preceded KRAB (Figure 4A, 4B, and S3B). For example, a 1-day overlap when KRAB stimulation preceded D3L led to ∼70% irreversibly silenced cells, whereas a 1-day overlap when D3L stimulation preceded KRAB only led to ∼30% irreversibly silenced cells (Figure 4B). In this case, it is likely that the 3-day prestimulation by KRAB, prior to overlap, allows more complete establishment of the silenced heterochromatin state, upon which D3L can more efficiently act. This result indicates that cooperation between KRAB and D3L is not merely due to co-occupancy of the factors at the target promoter, but rather that D3L only becomes highly effective at establishing memory after KRAB has completely silenced expression first.

Overall, these findings support and expand on a previous kinetic model of chromatin regulation, wherein KRAB and D3L promote the stepwise transition of a promoter first from active to transiently silenced and finally to the stably silenced state (Figure 4C).^40^ KRAB, probably through the establishment of H3K9me3 and HP1-dependent heterochromatin,^36^ catalyzes the transition from actively transcribed to transiently silenced. In contrast, D3L, probably through self-sustaining DNA methylation and the recruitment of additional chromatin silencers,^30^ catalyzes the transition from the *transiently* silenced state to the *stably* silenced state (Figure 4C). The notion that KRAB and D3L primarily function at distinct steps in this cascade further supports the finding that the synergy is only effective if the first transition, mediated by KRAB silencing, has been sufficiently made.

These findings further elaborate on the temporal relationship of these chromatin regulators,^53,66–71^ although the exact durations of each will probably vary in response to different input types and in different chromatin contexts. Nonetheless, these findings also show how the temporal relationship between these two forms of gene silencing can be exploited in synthetic circuits for discriminating input sequences.

### Combinatorial inputs can direct stable cell state transitions among alternative choices

Natural multi-potent cells can be guided to transition to one of several alternative states, based on the identity and combination of inputs.^72,73^ We reasoned that we could use our epigenetic memory switches to engineer a receiver cell that, similarly, can choose which of multiple distinct alternative states to transition to depending on the inputs it received. To achieve this, we expanded the initial circuit design (Figure 2A) by incorporating an additional epigenetically regulated promoter driving an mCherry reporter (instead of EGFP) as well as a new, third synNotch receptor that recognizes Her2 (Figure 5A).^74,75^ This mCherry promoter was targeted by this anti-Her2 synNotch receptor, which harbors an intracellular domain in which the LexA^76^ DNA binding domain is fused to the KRAB domain. In addition, the mCherry promoter also contains Gal4 operator sites that allow it to be targeted by the anti-nfGFP D3L synNotch receptor (Figure 5A). Thus, the receiver cell can adopt four possible states: EGFP^low^ mCherry^low^; EGFP^high^mCherry^low^, EGFP^low^mCherry^high^; and EGFP^high^ mCherry^high^ (Figure 5B). Correspondingly, this receiver cell can also sense combinations of three input signals, CD19, nfGFP, and Her2, presented on the surface of sender cells (Figure 5A).

**Figure 5.**
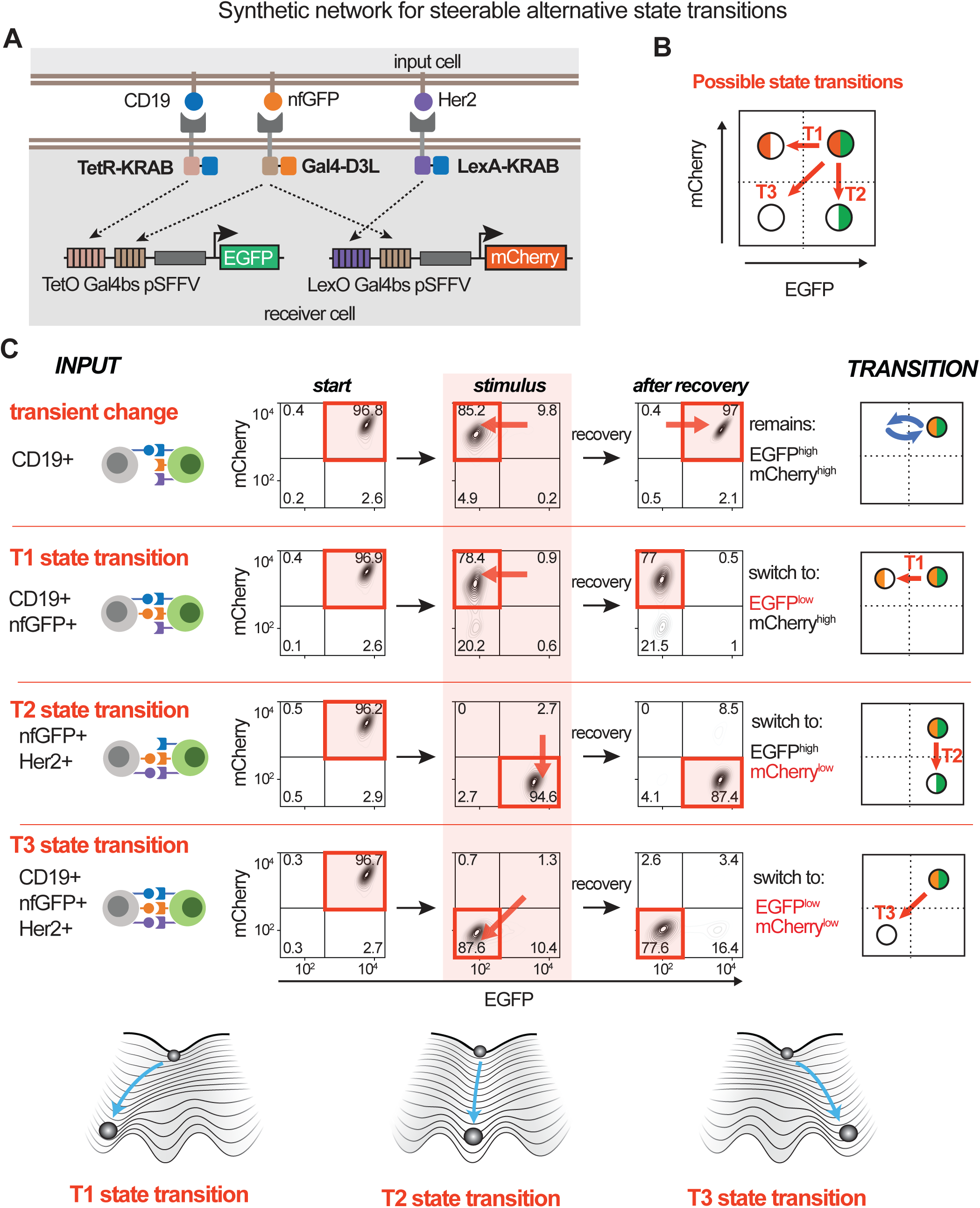
Input combinations can direct specific transitions from among alternative choices. **(A)** Design of a multipotent receiver cell type that can specifically silence either EGFP or mCherry depending on the combination of synNotch input signals. **(B)** Schematic illustrating the possible transitions of the multipotent receiver cell. Cells begin in the top right double-positive quadrant and can choose among three alternative transitions based on specific input combinations. **(C)** Two-dimensional flow cytometry distributions of EGFP and mCherry fluorescence. Receiver cells were co-cultured with different sender cells, as indicated in the ***INPUT*** column, for 7 days, and then allowed to recover in the presence of DAPT for 7 days. Numbers are the percentage of cells in each quadrant. Panels under ***TRAJECTORY*** summarize the state transitions. Distributions are representative of 3 technical replicates of a single cell clone. Below is a conceptual summary of how the inputs shape epigenetic landscapes to steer transitions to alternative states.

Altogether, this network design represents a minimal architecture that enables independent silencing of each target construct as well as independent control of both KRAB and D3L factors. Selection of the target to be silenced should be determined by the release of TetR-KRAB and LexA-KRAB synNotch factors, which occur upon CD19 and Her2 ligand stimulation, respectively (Figure 5A). However, the decision to make silencing irreversible depends on whether the cell receives the Gal4-D3L inducing signal, nfGFP. Here, a single Gal4-D3L module can be shared by both target promoters to determine commitment to silencing, since D3L does not substantially silence expression on its own. Consequently, the engineered cell, starting from the EGFP^high^ mCherry^high^ baseline state can, in principle, be directed to the three alternative states via distinct transitions, labelled as transitions T1 (EGFP^low^ mCherry^high^), T2 (EGFP^high^ mCherry^low^), and T3 (EGFP^low^ mCherry^low^) in Figure 5B.

As before, to avoid heterogeneity from reporters integrated at different local chromatin environments, we isolated three different isogenic clones harboring the complete network and examined each in parallel. We co-cultured the multipotent receiver cells with the different sender cell types for 7 days to ensure sufficient stimulation, followed by recovery in the presence of DAPT to assess committed cell state changes. As we had observed with the initial single target receiver (Figure 2), stimulation with CD19 alone, Her2 alone, or CD19 and Her2 together, which only induce recruitment of KRAB to the two target promoters, resulted in only transient changes to alternative expression states during stimulation (Figure 5C and S4A). However, co-culture with senders presenting the same ligand sets but also with nfGFP, which recruits D3L to both targets, resulted in transitions to each of the alternative stable states dictated by the specific KRAB synNotch variant that was stimulated (Figure 5C and S4B). CD19-nfGFP senders induced the T1 transition, Her2-nfGFP senders induced the T2 transition, and CD19-Her2-nfGFP senders induced the T3 transition, with little deviation to the other states (Figure 5C and 5D). As a further illustration of the modularity of the system, the T3 transition could be induced by co-culture with a single sender cell type expressing all three ligands, by two different sender cell types expressing CD19 and nfGFP or Her2 and nfGFP, or by three sender cell types each expressing CD19, Her2, or nfGFP (Figure S4C). Thus, the specific transitions between alternative states can be selectively controlled by the combination of stimulatory inputs.

We also realized that the irreversible commitment to each state would allow sequential transitions, in addition to the direct T3 transition, from the EGFP^high^ mCherry^high^ *baseline* state to the *terminal* EGFP^low^ mCherry^low^ state (Figure S4D). For example, a receiver cell experiencing Her2-nfGFP undergoes the T2 transition to the EGFP^high^ mCherry^low^ state. Exposure to a second stimulus after removal of the first, specifically CD19-nfGFP, should then result in a new transition (T5) from EGFP^high^ mCherry^low^ to the terminal EGFP^low^ mCherry^low^ state (Figure S4D). Indeed, we performed a stepwise co-culture program whereby we first co-cultured the multipotent receiver with Her2-nfGFP senders, removed the senders by FACS, and then exposed the resulting receivers to the CD19-nfGFP sender. Cells transitioned specifically between the states in the prescribed manner and remained in the terminal EGFP^low^ mCherry^low^ state after synNotch inhibition (Figure S4E). By comparison, performing stepwise input exposure without inclusion of D3L resulted in transient shifts between alternative states, but no ability to undertake multiple stepwise transitions toward a terminal state (Figure S4E). Thus, epigenetic memory can allow specific temporal sequences of transitions, where the history of inputs determines the order of transitions.

Altogether, the engineered multipotent receiver cell highlights the highly flexible and precise capabilities of the synNotch-based epigenetic switching system. This synthetic circuit forms a foundation for complex programmable differentiation systems.

### Stable gene activation switch using an inverter cascade

These types of irreversible epigenetic systems are able to effectively silence a target gene locus, but how can one irreversibly activate a gene locus? We reasoned that stable activation could be accomplished, without introducing additional chromatin regulators, by linking our modular silencing switch to a repressor that could act as an inverter module. More specifically, the epigenetic silencing of the repressor would relieve the repression of a second downstream gene, in turn leading to its activation (Figure 6A). To construct such a system, we first placed the bacterial transcriptional repressor protein, CymR,^77^ under the control of the engineered TetO-UAS-SFFV promoter, such that KRAB and D3L recruitment stably silences CymR expression.

**Figure 6.**
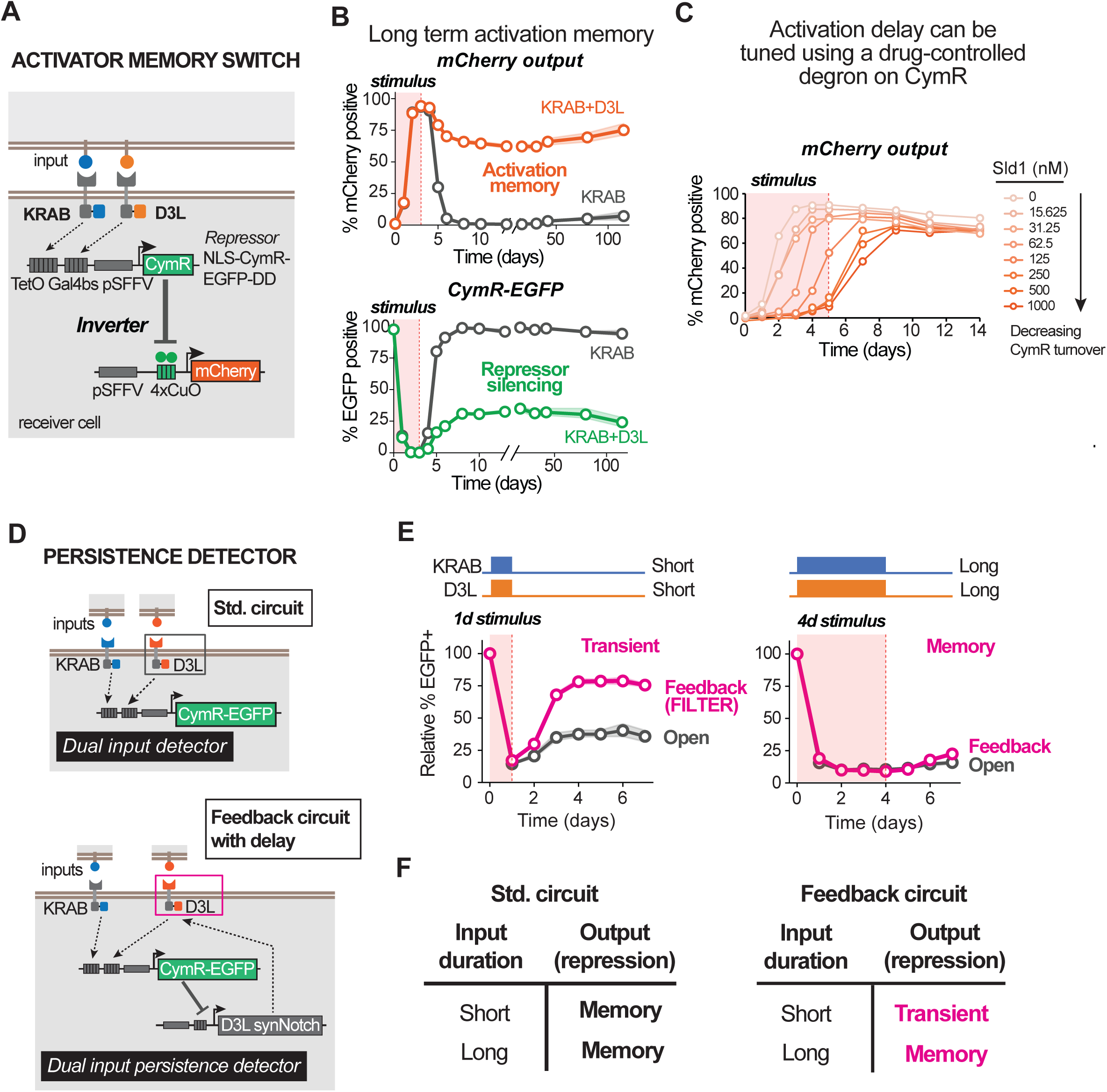
Engineering a stable gene activation memory switch using an inverter circuit. **(A)** Design of a receiver cell that incorporates an inverter cascade circuit architecture, whereby synNotch silencer receptors control an intermediate repressor, which in turn represses a downstream engineered target gene (mCherry). **(B)** Time courses showing the percent of cells expressing mCherry (top) or CymR-EGFP (bottom) following a transient stimulus (shaded area). Data for all curves are from 3 technical replicates of a receiver cell line derived from a single cell clone. Confidence intervals are the shaded areas surrounding the curves. **(C)** Time courses showing the percent of cells expressing mCherry following a transient stimulus (shaded area), and in the presence of varying amounts of the small molecule Sld1. Data for all curves are from 3 technical replicates. Confidence intervals are too small to be visible. **(D)** Design of a receiver cell in which the D3L synNotch is suppressed by CymR in the inverter cascade forming a feedback loop (bottom), compared to a standard (Std.) circuit design where both synNotch receptors are constitutively expressed. **(E)** Time courses showing the percent of cells expressing CymR-EGFP (relative to day 0) following 1-day or 4-day transient stimuli (shaded area). Data for all curves are from 3 technical replicates. Confidence intervals are the shaded areas surrounding the curves. **(F)** Truth table relating varying input durations with output responses based on the circuit design.

Second, we constructed a downstream locus expressing an mCherry reporter gene (with a PEST degron) that is directly inhibited by CymR, specifically by placing 4 copies of the CymR DNA binding site (CuO)^77^ immediately downstream of a separate copy of the SFFV promoter. This configuration results in inhibition of mCherry expression when CymR is expressed under baseline conditions (Figure 6A). CymR transcriptional inhibition should be readily reversible as it is mechanistically unlikely to lead to chromatin remodeling or epigenetic memory. Lastly, we also fused CymR to EGFP-DD, allowing us to monitor the state of this inverter cascade at both stages (expression of the repressor and of the output gene) following synNotch stimulation (Figure 6A). These constructs were integrated in different genomic locations via separate lentiviral vectors.

At baseline, cells express the CymR-EGFP gene, which represses mCherry expression from the downstream promoter (Figure 6B). However, upon KRAB and D3L synNotch stimulation, CymR-EGFP is quickly silenced followed by a matched upregulation of mCherry reporter expression (Figure 6B and S5A). When only KRAB is recruited, only transient mCherry expression is observed; inhibition of synNotch results in a loss of mCherry expression, corresponding to the reactivation of CymR-EGFP and restoration of baseline transcriptional repression (Figure 6B). In contrast, co-stimulation of both KRAB and D3L synNotch receptors resulted in long-term (≥100 cell generations) stabilized mCherry activation even after synNotch inhibition, consistent with epigenetic memory at the CymR-EGFP locus (Figure 6B). To further corroborate the strict inverse relationship between stable silencing of the CymR-EGFP locus and the stable activation of the mCherry output, we performed a series of stimulations of increasing duration from 1 to 3 days (Figure S5B). Similar to previous observations (Figure 3), shorter stimulus durations produced stable population bifurcation of the CymR-EGFP expression, and correspondingly, the mCherry output (Figure S5B and S5C). Importantly, bifurcation of CymR-EGFP was almost perfectly coupled to the bifurcation of mCherry, such that cells that silenced CymR-EGFP also completely activated mCherry and vice versa (Figure S5C).

Although we observed rigorous coupling between mCherry activation and CymR-EGFP silencing, the gene switches do not occur simultaneously. As previously observed with transcriptional cascades,^78^ upregulation of mCherry following synNotch stimulation occurs after a delay (Figure 6B and S5D), the duration of which can be explained by the time required for CymR-EGFP protein to sufficiently diminish, which in turn is dependent on its turnover rate. We reasoned, therefore, that the delay timing of mCherry activation can be tuned by modulating the stability of CymR-EGFP through the controllable degron, FKBP-DD (Figure 6A).^55^ Treatment of cells with increasing concentrations of the small molecule, Shield1 (Sld1)^79^ stabilizes the degron, resulting in slower CymR-EGFP downregulation following synNotch stimulation (Figure S5E).

Correspondingly, increasing Sld1 concentrations results in a greater delay time to activate mCherry (Figure 6C). Time to the first detectable increase in mCherry positive cells can be as short as 1 day and as long as 5 days following initiation of synNotch stimulation (Figure 6C). Thus, we can systematically tune the timing of this epigenetic activation circuit.

Lastly, we predicted that we could take advantage of this circuit activation delay to generate an input persistence detector. That is, we devised a system which requires longer stimulus durations to lock in epigenetic memory, and conversely, filters out short input pulses. To construct such a circuit, we placed the synNotch-D3L receptor under the control of the inverter activation switch. Under starting conditions, the D3L synNotch receptor is transcriptionally repressed by CymR-EGFP (Figure 6D). In turn, when both synNotch-KRAB and synNotch-D3L ligands are presented but for only a short duration (∼1 day), receiver cells initiate silencing of the CymR-EGFP locus, but only transiently (Figure 6E and 6F), as only the KRAB synNotch input is detected. Because the D3L synNotch is repressed initially, the cell essentially ignores the D3L input signal. However, if the dual input persists long enough for CymR to fully turn over, then the synNotch-D3L receptor can be expressed. In turn, a longer persistent input (∼4 days) can give cells time to sense both KRAB and D3L inputs, allowing them to establish epigenetic memory and stably silence CymR-EGFP (Figure 6E and 6F). Thus, only an input pulse long enough to overcome the delay of CymR turnover results in epigenetic memory.

### Epigenetic switching can yield stable morphogenic changes

We next sought to demonstrate if these epigenetic state switching circuits could be used to induce functionally important irreversible changes in cell behavior. We focused on the ability to alter a cell’s physical adhesion and sorting properties, a type of state change that plays a central role in development and other multi-cellular self-organizational processes. In development, cell adhesion plays a crucial role in the formation of distinct spatial domains and morphogenic structures.^80,81^ Changes in cell adhesion, such as the expression of cadherin proteins, can induce a population of cells to form compact aggregates due to strong homotypic interaction.^80^ Within a mixed multicellular population consisting of cells expressing differing types of cadherins, distinct cell populations will differentially segregate from one another into distinct domains.^81–84^ This type of differential adhesion sorting^85^ is used throughout development, resulting in downstream cell lineages that move or migrate to new positions to further elaborate morphological structures in the body. Critically, for this kind of spatial sorting to be stable over the course of development, changes in expression of the responsible adhesion molecules must be irreversible.

We first designed a circuit in which interaction of a sender cell with a receiver cell would induce expression of the adhesion molecule, P-cadherin (PCAD). We linked PCAD to the epigenetic activation switch described above. Specifically, we designed an mScarlet-P2A-PCAD fusion protein that is controlled by the CymR-EGFP-DD protein (Figure 7A). Following synNotch stimulation of both KRAB and D3L, mScarlet and PCAD expression can be stably induced (Figure S6A). We confirmed that co-culture of receiver cells with the proper sender cells would induce PCAD-driven compact aggregation and self-organization (Figure 7A and 7B). Importantly, when cells are co-cultured in an ultra-low attachment (ULA) culture plate, the differential adhesion properties between the sender cells that do not express a cadherin and the activated receiver cells expressing PCAD results in the morphogenesis of a compact core of receivers surrounded by an outer shell of sender cells (Figure S6B). This confirms the ability to switch between cell adhesion phenotypes by way of a synNotch-gated inverter cascade circuit.^83^

**Figure 7.**
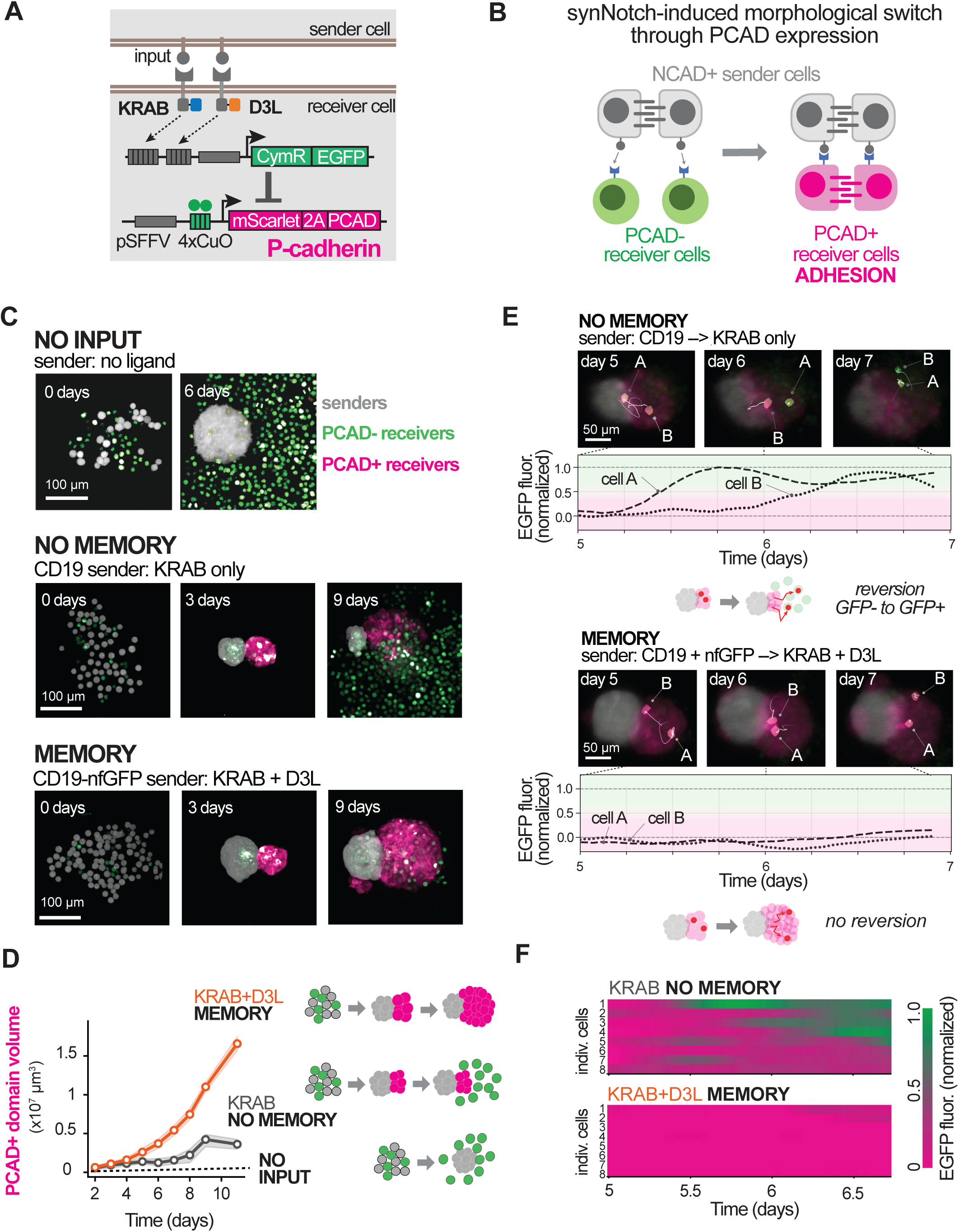
Epigenetic memory stabilizes morphogenesis in a dynamic self-organizing system. **(A)** Design of a receiver cell that activates PCAD expression in the inverter cascade. Note that the cells also constitutively express a synthetic adhesion molecule (synCAM), which we have found to disrupt baseline cell-cell adhesion. This synCAM does not participate in the epigenetic switching circuit that controls P-cadherin and is not illustrated for simplicity. **(B)** Schematic highlighting the outcome of circuit activation in receiver cells upon contact with sender cells. Receiver cells undergo a morphological change as PCAD mediates strong homotypic adhesion. Additionally, as the sender cells also express a different cadherin, NCAD, sender and activated receiver cells can self-organize into distinct spatial domains. **(C)** Confocal microscopy images (maximum intensity projections) for an experiment where senders and receivers were co-cultured in an ultra-low attachment vessel. Images for NO MEMORY and MEMORY sets are representative of 5 technical replicates and of 3 single cell clones (see Figure S6D and S6E). **(D)** Time course of the size of the PCAD+ mScarlet+ domain during the self-organizing dual spheroid experiment. 3D Volumes were measured from the confocal microscopy experiment shown in (C) and replicates shown in Figure S6D. The cartoons to the right summarize the morphological dynamics. **(E)** Light sheet microscopy images and corresponding time courses of normalized EGFP fluorescence intensity of two example individual receiver cells tracked within the induced spheroid structures. The thin white lines in the microscopy images correspond to the trajectory of the labeled cells from the prior 6 hrs. The trajectories of these single cells are shown in Video S1. **(F)** Heatmaps showing time courses of normalized EGFP fluorescence intensity for a panel of tracked cells within the induced spheroid structures.

We then tested whether epigenetic control of PCAD adhesion could lead to irreversible morphological changes, even when differential sorting of the PCAD+ receiver cells causes them to move out of contact with the sender cells that originally induced the change. This situation would mimic many situations in development, in which an inherently transient interaction induces changes in a cell that must be maintained, even after it has migrated or sorted away from the inducing signals. We designed a co-culture experiment in which we used sender cells expressing N-Cadherin (NCAD) and receiver cells that do not express a cadherin at baseline, but that induce the expression of PCAD upon receiving synNotch stimulation. Importantly, NCAD and PCAD cells will differentially sort from one another (because PCAD-PCAD and NCAD-NCAD homotypic interactions are stronger than PCAD-NCAD heterotypic interactions).^86^ Thus, when receiver cells induce PCAD expression, they should segregate from the NCAD sender cells (Figure S6C), creating a situation in which the sender-receiver interaction is inherently transient. Thus, the receiver cells would only maintain their newly induced adhesion state if they irreversibly activated PCAD expression.

We constructed three alternative types of NCAD expressing sender cells. Specifically, we generated control cells that do not express a synNotch ligand (negative control), sender cells that express the synNotch-KRAB ligand (CD19 – no memory), or senders that express both the synNotch-KRAB and synNotch-D3L ligands (CD19 and nfGFP - memory). Thus, each of these populations of NCAD sender cells invoke a different response in the receiver cells. To simplify these co-culture experiments, we pre-treated sender cells with mitomycin C to arrest their proliferation. This way, sender cells can continue to stimulate receivers throughout the experiment, but the overall sender cell population does not substantially grow in size thereby maintaining a limited sender-receiver interface area.

When receiver cells were co-cultured with the control cells with no ligand, the receivers remain in an EGFP^high^ state and do not aggregate (Figure 7C). When the same receivers were mixed with the CD19+ senders that stimulate synNotch-KRAB only, CymR-EGFP was largely silenced and cells induced mScarlet expression (and linked PCAD), leading to self-organization into a PCAD-based aggregate adjacent to the sender cell aggregate (Figure 7C, S6D, and S6E). Importantly, as the receiver cells continue to grow, and because the interfacial region between the sender and receiver cells is finite, the receiver cells lose contact with the senders and we correspondingly observed a large reversion of the receiver cells to the PCAD- unclustered adhesion state (EGFP expression returns, and the cells dissociate) (Figure 7C, S6D, and S6E). Only a subpopulation of receiver cells that remain in close contact with the sender cells were able to maintain the morphogenic switch.

In contrast, when receiver cells are cultured with the CD19+ nfGFP+ sender cells, which stimulate both KRAB and D3L and result in memory (Figure S6A), we again observed largely irreversible changes in adhesion and EGFP expression. Receiver cells induced mScarlet and PCAD within a few days, resulting in a self-organizing aggregated cluster adjacent to the sender cell mass (Figure 7C, S6D, and S6E). Crucially, in the presence of this sender cell type, most of the receiver cells remain in the mScarlet+ PCAD+ state, even as the spheroids grow and the outer regions lose direct juxtacrine input from the senders (Figure 7C, S6D, and S6E). Using confocal microscopy, we measured the volume of the resulting PCAD+ domains over time and found that cells that received KRAB+D3L input signals produced a spheroid that continuously grew, becoming substantially larger than the KRAB-only induced spheroids (Figure 7D). These observations are consistent with the KRAB+D3L input enabling the irreversible maintenance of the PCAD+ state over time and space, whereas KRAB input alone induced only an unstable reversible population of PCAD+ cells.

To more closely analyze these morphogenic outcomes at the individual cell level, we sought to trace individual receiver cell states over time. We predicted that we would observe individual induced EGFP- PCAD+ cells reverting to the baseline EGFP+PCAD- state over time with KRAB-only input, thereby reflecting the global instability of the induced spheroid. To do this, we sparsely labeled a small fraction (1/10) of receiver cells using a nuclear-localized iRFP670 label, which enabled us to track the individual cells within the assembly by timelapse light-sheet microscopy. We used the nuclear marker to define individual cells, trace their trajectories, and measure the fluorescence intensity within the cell envelope over several days. Receiver cells dynamically and stochastically move within the receiver aggregates with some cells appearing to recontact the sender aggregate whereas others move away.

With KRAB-only stimulation we observed clear examples of individual induced receiver cells reverting from the EGFP-PCAD+ state back to the baseline EGFP+PCAD- state (Figure 7E, 7F, and Movie S1). However, with KRAB+D3L stimulation, which induced memory, we could not observe any such receiver cell reversion events, even though cells also stochastically move within the receiver aggregate and away from the sender interface. This behavior of the KRAB+D3L stimulated receiver cells is consistent with an irreversible commitment to the PCAD+ state following transient sender cell contact (Figure 7E, 7F, and Movie S1). Individual cell states that we tracked over time are summarized in Figure 7F and S6G.

Altogether, these results demonstrate that the epigenetic switching circuits can be readily adapted to control transitions between different cell adhesion states. In turn, we have shown how irreversibly inducing a stable morphogenic change, such as a change in adhesion properties, unlocks the potential for more complex trajectories of multicellular self-organization, especially ones that allow memory of a history of past transient inputs.

### Epigenetic memory circuit allows short input pulses to drive fibroblast reprogramming toward a neuron-like state

To examine the potential of these epigenetic memory circuits to modulating the endogenous transcriptome and natural cell states, we sought to test whether it could be used to drive the reprogramming of fibroblasts toward a neuron-like state. Such a process is challenging due to the requirement for sustained expression of combinations of lineage-specifying transcription factors (e.g., Brn2, Ascl1, and Myt1l = “BAM” factors)^87^ that substantially remodel gene regulatory networks and the epigenetic landscape.^88,89^ Conventional transdifferentiation protocols entail prolonged induced expression of these factors, often over periods of weeks, to achieve appreciable conversion.^87,88,90,91^

We reasoned that a synNotch-based memory circuit capable of inducing prolonged transcription factor expression in response to a transient, brief initiating stimulus could drive this lengthy process. In other words, our memory circuit would allow cells to commit to undergo reprogramming. More broadly, a receptor-mediated state conversion would be a foundation for advanced tissue engineering with spatial and temporal control. Therefore, we deployed the epigenetic inverter cascade to drive expression of the neuronal reprogramming BAM factors in C3H immortalized mouse embryonic fibroblasts (Figure 8A).^87^ Cells were transduced with lentiviruses encoding the five components of the circuit, enabling for synNotch-responsive activation of BAM expression. As a control non-epigenetic induction system, we also generated a C3H cell line in which the BAM factors could be induced by doxycycline (Figure 8A).^87,92^ For each induction system, we could then test whether a short or long input stimulation was required to commit to a neuron-like conversion, as assessed by detection of the neuronal marker, Class III β-tubulin (marked by the Tuj1 antibody) and observing the formation of thin neurite-like processes by microscopy (Figure 8B-D, and S7B). In both cases, cells were stimulated for either 2 days (short input pulse) or 10 days (long constant input), and the number of cells displaying the neuron-like cell state was assessed periodically for up to 10 days.

**Figure 8.**
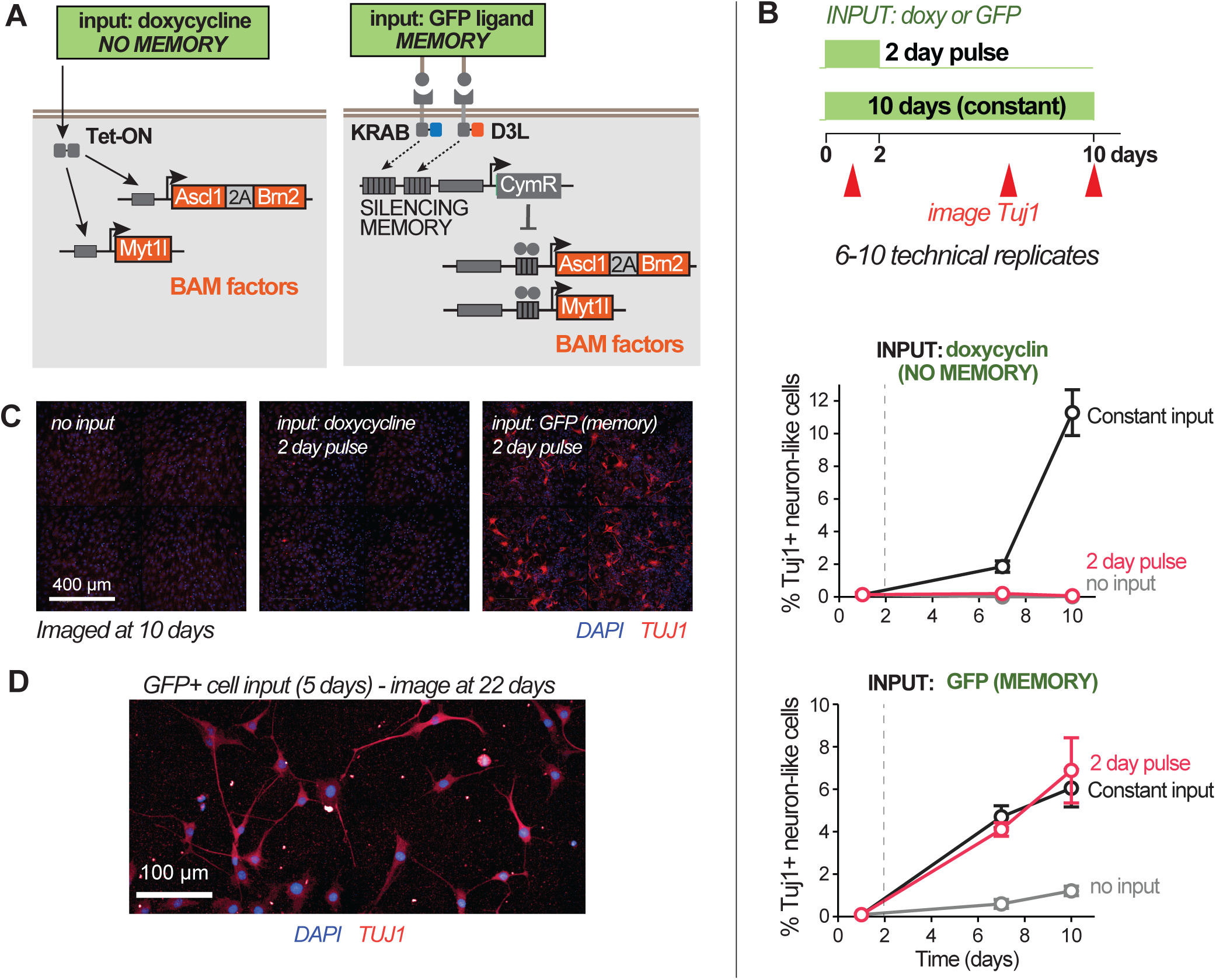
Epigenetic memory circuit enables a short input pulse to drive fibroblast reprogramming toward a neuron-like state. **(A)** Design of cells that activate Brn2, Ascl1, and Myt1l (BAM factors) expression via either doxycycline-inducible Tet-On (left) or the synNotch-KRAB-D3L inverter cascade (right). Note that memory circuit cells also express an mCherry reporter together with the BAM factors but this is not shown for simplicity. Additionally, this receiver cell harbors two synNotch receptors that contain KRAB and D3L regulators, but both recognize GFP as a ligand. **(B)** Time courses of the percent of observed Tuj1+ neuron-like cells relative to the total number of C3H cells initially plated. C3H cells harboring the constructs shown in (A) were transiently stimulated for 2 days or constantly for 10 days (green bars) by surface-immobilized GFP ligand. Values shown are average of 6-10 technical replicates with error bars depicting standard error. **(C)** Confocal microscopy images showing cells stained with Tuj1 antibody at 10 days after the initiation of BAM factor expression, and in response to no input or 2-day input pulse of doxycycline or GFP ligand. **(D)** Same as (C) but where cells were stimulated in a co-culture with L929 fibroblast sender cells presenting the GFP ligand. Note that L929 senders express iCaspase9 and were induced to undergo apoptosis at day 5 of the co-culture in turn leading to a 5-day input pulse. Visible in the microscopy field are the remnants of the senders (green).

We first examined non-epigenetically induced expression of BAM factors (via doxycycline). When these cells were stimulated for ten days continuously, we observed a significant number of Tuj1+ neuron-like cells (Figure 8B and 8C). However, the short stimulus pulse of 2 days failed to produce neuron-like cells after 10 days of incubation (Figure 8B and 8C). These data indicate that a relatively long input duration and sustained BAM expression was required for cells to commit to the reprogramming process. We then induced activation of the epigenetic circuit by surface-immobilized GFP ligand. Upon stimulation, we observed stable induction of the mCherry reporter, indicating BAM factor expression (Figure S7A). In contrast to the doxycycline-based BAM induction, we observed that the short 2-day pulse of synNotch-KRAB-D3L stimulation, via GFP ligand, led to a substantial number of cells acquiring a neuron-like state by day 10. The reprogramming rate was essentially indistinguishable from that observed with 10 days of constant GFP ligand input (Figure 8B and 8C). We also tested whether transient cell–cell contact with GFP ligand-presenting sender cells could induce reprogramming and indeed found that a 5-day pulse was also sufficient to produce Tuj1+ neuron-like cells (Figure 8D). We further characterized the cells induced by a 5-day GFP ligand pulse, and found that they were positive for additional neuronal markers, specifically vesicular glutamate transporter, vGlut1, and glutamate decarboxylase, Gad65/67 (Fig S7B).^87^ These markers, associated with excitatory and inhibitory neurons, respectively, have both been observed in published BAM-mediated neuronal reprogramming studies.^87^ The presence of the these additional markers further support that cells were indeed reprogrammed away from a fibroblast state toward a neuron-like state in response to the transient GFP ligand-to-synNotch input.

In summary, these findings demonstrate that the synthetic epigenetic memory circuit can be leveraged to produce long lasting changes in the expression of endogenous genes, in this case driving early commitment to reprogramming. In general, this epigenetic circuit-based reprogramming approach should offer a route to efficient generation of target cell types, particularly in contexts where input signals may be of limited duration, and where spatial control over the reprogramming event is required.

## DISCUSSION

### Engineering modular input-induced epigenetic cell state switches

In development, long-term commitment to differentiated cell fates is controlled by the convergence of cell signaling with epigenetic gene regulatory systems.^93,94^. In this work, we developed a series of synthetic circuits that harness the epigenetic regulatory machinery to drive user-specified signal-induced cell state transitions. These modules represent a generalizable and customizable approach to control transitions to specific stable cell states. Here, we consider how irreversible changes in gene expression can be a foundation for controlling cell states. Our synthetic epigenetic circuits provide a means to dynamically tune bifurcation leading to population diversification. Furthermore, these synthetic modules can be used in combination. They can be scaled up to regulate multiple genes independently and selectively, and they can also be linked in series to drive a specific trajectory of state changes. Given that this approach relies on harnessing endogenous epigenetic regulatory mechanisms, these types of synthetic circuits could be applicable in diverse mammalian cell types.

### Alternative approaches for engineering multiple stable cell states

This work represents one of the first examples of incorporating epigenetic regulators KRAB and Dnmt3 in synthetic gene networks that drive selective and irreversible cell state switches. Previous efforts uncovering the potent synergy of KRAB and Dnmt3 chromatin mechanisms for long-term stable silencing have primarily focused on using this multi-factor approach to silence endogenous genes for gene therapy applications.^24,25,45,46,54^

Prior studies in synthetic biology have explored alternative approaches to achieve irreversible or stable cell state switches. One approach is the use of networks of mutually inhibitory transcription factors to create bistable and multistable systems.^15–17^ Another is to use recombinase-triggered systems to flip selective promoter-gene relationships yielding stable inhibition or activation.^95–98^ Others have used orthogonal systems that mimic the writer-reader functionality of natural epigenetic systems.^99^ While each of these approaches has its advantages and highlights different principles, our approach of recycling the native epigenetic machinery (and the power of DNA methylation transmission as a source of memory) is highly modular, and therefore lends itself to constructing higher order systems in which transitions are highly targeted, tightly controlled by specific combinations and durations of signals, and can be linked together into specific transition cascades. Moreover, all of these diverse synthetic memory systems have the potential to be linked together to create hierarchical systems that integrate different mechanisms of cell state determination and stability.

### Cooperativity between KRAB and Dnmt3 in stable silencing

The switches we have built depend on the cooperation between the KRAB domain (and the KAP1 complex that it recruits) with the Dnmt3 complex to reconfigure chromatin to generate a self-sustaining transcriptionally inactive state. The combinatorial action of these factors—one that drives heterochromatin formation and one that drives DNA methylation—has also been harnessed recently as a means of targeting endogenous genes for long-term silencing by fusing KRAB, Dnmt3A, and Dnmt3L factors to dCas9.^24,25,45^. Indeed, many studies have identified several mechanisms of interplay between heterochromatin formation and DNA methylation that can account for this cooperativity. Such potential mechanisms include direct interactions between heterochromatin protein 1 and the Dnmt3 complex,^66,67^ potentially enhancing its localization when KRAB/KAP1 establishes a silenced heterochromatin state. Additionally, it is well established that histone H3 lysine 4 (H3K4) trimethylation, which is strongly associated with active transcriptional activity at promoters, does not allow full activity of Dnmt3A.^68,69^ Thus, silencing of transcription, which is associated with H3K4 demethylation, may produce the chromatin environment allowing Dnmt3 to efficiently methylate DNA. In support of this, it was recently found that fusing Dnmt3L to the H3 N-terminal domain, which effectively provides an unmethylated H3K4 signal, potentiates DNA methylation and silencing by the Dnmt3 complex.^54^

Our findings are consistent with these proposed mechanisms, and provide critical insight into the temporal parameters required for effective cooperation of these chromatin factors. By systematically altering stimulation duration and relative timing of KRAB and D3L recruitment, we provide support for a simplified model describing the transition to a silenced state as a two-step process. Our findings are consistent with the Dnmt3 complex participating in the second step of this process, that is, being the primary driver of establishing the *stably* silenced state from the *transiently* silenced state. This explains why D3L is largely ineffective at establishing a silenced state on its own (i.e., without KRAB). In contrast, KRAB appears to primarily impact the first step, to potently overcome active transcription and in effect, prime D3L for its function by producing a permissive chromatin state for it to act. Of course, this model depends on the chromatin context at an ectopic constitutive promoter, such as the SFFV promoter, and differences probably arise in different genomic regions. It would be interesting for future analyses to consider how different classes of promoter or enhancer sequences might alter these dynamical behaviors.

### Controlling cell state trajectories over time and space

Inducible and programmable epigenetic silencing can be an important foundation for engineering cell state and fate switches. In future cell engineering applications, synthetic memory circuits would be most valuable when signals are inherently transient and/or evolve over time, as in the case of development-like processes with morphogenesis and differentiation. Note that in this work, we have used DAPT as an experimental tool to artificially produce transient signals and demonstrate memory of a gene expression switch. However, DAPT is not itself a foundation of the epigenetic system nor is it required to drive any fate reprogramming. In this work, we examined how epigenetic memory can regulate cell adhesion—a demonstrative example in which the induced state change itself drives the dynamic evolution of the system, wherein responding cells become segregated from the source of stimulus. Effectively, cells experience a transient signal, (without the addition of DAPT) because the induced state change segregates activated receiver cells from the sender cells. Nonetheless, we clearly observed that such an inherently transient cell-cell interactions in this system can lead to an irreversible cell state change.

As the complexity of multicellular engineering increases, such tools could give us the foundation to engineer differentiation trajectories determined by sequences of inputs separated in space and time. We also show that these epigenetic switches can be used to reprogram endogenous gene programs, such as inducing fibroblasts to enter a neuron-like state in response to a transient signal. Enabling cells to flexibly decide whether to transiently change their expression or to commit to a new state, raises the possibility of engineering far more dynamic and complex cellular systems.

Circuits that harness epigenetic memory may be useful for engineering synthetic development and regeneration processes that require complex multi-stage stepwise differentiation.^100^ We envision circuits that could enforce a sequence of state transitions where order matters for final function.^101^ Engineered cells would only commit to differentiate when they sense the right combination and timing of input signals. The key premise is that memory and state commitment are fundamental behaviors that allow cells to integrate multiple inputs over time and space, and likewise program transitions in time and space.

Similarly, epigenetic circuits could be deployed to engineer cell state bifurcation, where a multipotent precursor cell may be programmed to diversify into different complementary cell types. One fundamental challenge for next generation cell therapies is the necessity to perform multiple cell functions simultaneously. For example, can we mitigate inflammation while simultaneously promoting tissue regeneration^102^ in inflammatory disorders? Instead of developing multiple cell products for each role or cell type, one could generate a single engineered cell population and use input-controlled epigenetic memory circuits to autonomously bifurcate the cell population into two or more cooperating states.

Broadly, we believe developing synthetic circuits capable of programming committed state transitions opens the door to cell-based technologies that can act in ways that parallel natural developmental processes, but with the control and precision of engineering.

## LIMITATIONS OF THE STUDY

This study uses lentivirus to transduce constructs into mouse fibroblasts, which generates variants in which constructs are integrated semi-randomly at sites throughout the genome. Specific loci contexts may differ significantly, with some regions either more primed or more resistant to KRAB or D3L function. Indeed, similar chromatin context effects have been recently noticed to impact prime editing.^103^ This loci-specific heterogeneity may explain why, for example, in some cases D3L stimulation alone leads to silencing in a small sub-fraction of the population. To address this, we have isolated multiple single cell clones of each circuit to allow us to study circuit behavior in a homogeneous context, but we appreciate that behavior at different loci with different base promoters or even in different cell types may be different. Future studies would ideally focus on targeting these systems to different specific loci with different properties, as well as incorporate additional chromatin regulatory components that might alter each locus’ propensity for epigenetic silencing.

Additionally, it is possible that the current iteration of the epigenetic circuit may require modification and optimization for use in other cell types. For example, pluripotent stem cells display higher DNA demethylation activity,^104–106^ which likely would interfere with the Dnmt3A/L mechanism our current circuits harness. In any case, the goal in this manuscript is to establish a conceptual foundation for approaches in which chromatin-based regulation can be harnessed into circuits for various spatiotemporal behaviors (memory, filters, alternative choices, etc). We fully expect that these types of capabilities will be possible to achieve in essentially any cell type, while acknowledging that specific components would need to be tailored to account for the cell-specific endogenous chromatin machinery.

Lastly, we acknowledge that our current implementation of reprogramming fibroblasts toward a neuron-like state does not produce more or better neurons than existing constitutive lentiviral fibroblast-to-neuron protocols. Also, our use of immortalized C3H fibroblasts that harbor multiple lentiviral integrations could be an obstacle for generalizing this control approach to primary cells. However, the innovation of this study is to show that a receptor-triggered circuit can drive cells to commit to a reprogramming process even with relatively brief extracellular inputs. More broadly, our work provides a first of its kind demonstration of how spatiotemporal cell-cell communication can be engineered to drive specific cell state reprogramming.

## ACKNOWLEDGEMENTS

This work was supported by the following: NIH/NIDA R01 DA036858 (W.A.L), Howard Hughes Medical Institute (W.A.L.), NSF DBI-1548297 (W.A.L.), NIH/NCI R01 CA253017 (W.A.L., R.A.), Helen Hay Whitney Foundation (O.A.C.) Cancer Research Institute (A.M.), and Additional Ventures Catalyst to Independence Award (Z.L.G.). Research reported in this publication was supported by the National Cancer Institute of the National Institutes of Health under Award Number R01CA253017. The content is solely the responsibility of the authors and does not necessarily represent the official views of the National Institutes of Health. We thank Geeta Narlikar, Wesley McKeithan, Levi Rupp and members of the Lim laboratory and UCSF Cell Design Institute for discussions, assistance, and advice. We thank the UCSF Center for Advanced Light Microscopy for light-sheet microscopy services.

## AUTHOR CONTRIBUTIONS

Conceptualization, W.A.L., R.A., and O.A.C; Formal analysis, O.A.C., A.M., S.R.W., Z.C., and Z.L.G.; Funding acquisition, W.A.L., and R.A.; Investigation, O.A.C., A.M., S.R.W., Z.C., Z.L.G., P.L., and R.A.; Methodology, O.A.C., A.M., S.R.W., Z.L.G., S.T., and R.A.; Resources, W.A.L., and B.G.B.; Supervision, W.A.L., R.A., and B.G.B.; Visualization, O.A.C., A.M., S.R.W., Z.C., Z.L.G., and R.A.; Writing – original draft, O.A.C., and R.A.; Writing – review & editing, O.A.C., R.A., W.A.L., A.M., Z.L.G., and S.T.

## DECLARATION OF GENERATIVE AI AND AI-ASSISTED TECHNOLOGIES

During the preparation of this work, the authors used Microsoft Copilot, Perplexity, and Claude to assist with generating Python code for microscopy analysis, figure plotting, and some initial text drafting. After using these tools, the authors thoroughly reviewed and edited the code and text for accuracy. The authors take full responsibility for the content of the publication.

## STAR METHODS

### Resource availability Lead contact

Further information and requests for resources and reagents should be directed to the lead contact Wendell Lim. To ensure a fast response, please copy Noleine Blizzard and Pilar Lopez in any requests related to the paper.

### Materials availability

Plasmids will be distributed via Addgene. Cell line requests can be made to the lead contact.

### Data and code availability

- Raw and analyzed data associated with this paper will also be shared by the lead contact upon request.
- Original codes used for processing data and plotting figures will also be shared by the lead contact upon request.
- Any additional information required to reanalyze the data reported in this paper is available from the lead contact upon request.

### Experimental Model and Study Participant Details Cell lines

All cells were handled in sterile safety cabinets, and cell densities were determined using the Countess 2 automated cell counter (Thermo Fisher). Mouse L929 cells (male) were cultured in Dulbecco’s Modified Eagle’s Medium (Thermo Fisher) supplemented with GlutaMAX and 10% fetal bovine serum (UCSF) at 37 °C with 5% CO_2_ in humidified incubators. Mouse C3H/10T1/2 Clone 8 (C3H) cells were cultured in Dulbecco’s Modified Eagle’s Medium (Thermo Fisher) supplemented with GlutaMAX, 1 mM sodium pyruvate (Thermo Fisher), 1X MEM non-essential amino acids (Fisher Scientific) and 10% fetal bovine serum (Thermo Scientific) at 37 °C with 5% CO_2_ in humidified incubators. They were maintained at a confluence approximately between 10% and 90%. For routine passaging, we lifted cells using TrypLE Express (Thermo Fisher). However, for modified L929 cells expressing the LaG16-LFA1β2 gene,^107^ which diminishes cell adhesion, cells were simply resuspended by vigorous pipetting as trypsinization is not required. Lenti-X HEK293T cells (female) were treated similar to L929 and C3H cells. All cell lines were used for experiments before they reached 10 passages, which were counted starting from the time the cells were frozen down to make stocks.

### Method Details

#### Study Design

The goal of this work was to demonstrate how modular genetic circuits incorporating chromatin regulators could be deployed for the engineering of cell state transitions. We iterated through varying designs to explore different features of the cell circuits, including DNA binding domains and synNotch receptors. We used cell and molecular biology experiments to assess synthetic circuit performance, including flow cytometry, cytosine methylation sequencing, and fluorescence microscopy. Results shown represent at least three replicates, as indicated in figure legends. This study was not blinded.

### Plasmid design and construction

SynNotch receptors harboring the chromatin regulators were constructed by iterating through the various domains. First, all synNotch variants were constitutively expressed by the human Elongation Factor 1-alpha promoter (EF-1α). All receptors contain an N-terminal CD8α signal peptide for membrane targeting and c-Myc, HA, or 3xFlag epitope tags. The epitope tags served a dual purpose for determination of surface expression and for receptor stimulation with magnetic beads. Recognition domains consisted of a the anti-CD19 scFv,^108^ anti-GFP LaG16 nanobody,^50^ or the anti-HER2 scFv clone 4D5-8.^74^ All receptors contained the mouse Notch1 minimal regulatory region (Ile1427 to Arg1752). The intracellular domains consisted of the yeast Gal4 DNA binding domain, the TetR DNA binding domain, the CymR DNA binding domain^77^ with an N-terminal 2-copy SV40 nuclear localization sequence (NLS), or the LexA DNA binding domain with an N-terminal c-Myc NLS. Lastly, the C-terminal end of the receptors consisted of the Kox1 KRAB domain^38^ or the mouse full-length Dnmt3L. All chromatin regulator components were also separated from the DNA binding domain by a short linker.

The target loci of the chromatin regulators were cloned into separate vectors. At the 5’ end, the constructs consisted of 7 copies of the tetracycline responsive element, followed by 8 copies of the ZFHD1 binding sites (present here only as a spacer), and lastly 5 copies of the Gal4 DNA binding domain target sequence. An alternative construct used for characterizing KRAB-D3L temporal overlap on memory establishment (Figure 4) consisted of 6 copies of the CymR targeting sequence instead of the Gal4 binding site array. Another construct used for the multi-state receiver cell (Figure 5) consisted of 6 copies of the LexA DNA binding domain target sequence followed directly with the 5 copies of the Gal4 DNA binding domain target sequence, as above. All binding site arrays were followed by the constitutively active SFFV promoter, which then drove expression of various transgenes, including EGFP, mCherry, or the CymR DNA binding domain. Note that the CymR domain was directly fused to EGFP and 2 copies of the SV40 NLS. All targets were in turn fused at the C-terminal end to an FKBP12 destabilization domain that is regulated by a small molecule, Shield1.^79^

The target loci of CymR^77^ were also cloned into separate vectors with redesigned promoter regions. The 5’ end consisted of the same SFFV promoter, but without the upstream arrays of binding sites. Instead, the SFFV promoter sequence was followed immediately downstream by 4 copies of the CymR target sequence. This promoter construct drove expression of mCherry fused to a PEST degron,^109^ or to mScarlet fused to an FKBP12 destabilization domain. For constructs used to induce morphogenesis (Figure 7) mCherry or mScarlet fragments were fused to a self-cleaving P2A peptide followed immediately by the wild-type mouse P-cadherin. For constructs used to induce fibroblast reprogramming (Figure 8) an mCherry-P2A fragment was fused to either a Brn2-P2A-Ascl1 chain (mouse genes) or to the mouse Myt1L gene. These vectors also included a separate downstream transcription unit consisting of the mouse phosphoglycerate kinase (PGK) promoter constitutively driving expression of either TagBFP or cell surface ligand consisting of the ALFA tag^110^ fused to a fibcon domain^111^ and finally the Intercellular Adhesion Molecule 1 (ICAM1) intracellular domain (ICD).^107^ Constitutive TagBFP and ALFA-fibcon-ICAM were used as selectable markers for FACS sorting.

Other constitutively expressed genes were also cloned into separate vectors. SynNotch ligands were constructed by fusing the CD19 extracellular domain, GFP, non-fluorescent GFP mutant protein,^56^, or the Her2 extracellular domain to the ICAM1-ICD.^107^ Similarly, an ALFA-fibcon construct was fused to the beta2 integrin ICD from the Lymphocyte function-associated antigen-1 (LFA-1) gene.^107^ Ligand constructs also harbored an N-terminal signal peptide for membrane targeting and either Flag or HA epitope tags for determination of surface expression via antibody staining. Ligands were constitutively expressed by the SFFV promoter. iRFP670 constructs for sender cells included the EF-1α promoter to drive constitutive expression in sender cells and additionally contained an N-terminal 2xSV40-NLS motif for nuclear labeling of receiver cells in self-organizing morphogenesis experiments (Figure 7). Mouse N-cadherin was expressed together with either iRFP670 or TagBFP in the form of an NCAD-P2A-iRFP670/TagBFP construct. The catalytic domain of the human TET1 gene was fused to the TetR DNA binding domain and expressed from the SFFV promoter. This construct also harbored an IRES element linked to TagBFP as a marker for FACS sorting. Lastly, an iCaspase9-P2A-mNeonGreen^112,113^ was expressed from the EF-1α promoter.

All of the constructs were cloned into a pHR’SIN vector via In-Fusion (Takara Bio). Inserts were PCR amplified using either the KAPA HiFi DNA Polymerase (Roche) or the Phusion High-Fidelity DNA Polymerase (New England Biolabs) with primers containing complementary ends for In-Fusion assembly. In other cases, double stranded DNA fragments were designed and ordered from Twist Bioscience and used directly for In-Fusion assembly. Vectors were linearized by restriction digests. PCR-produced inserts and digested vectors were extracted from agarose gels using the Zymoclean Gel DNA Recovery Kit (Zymo Research) and used for In-Fusion reactions. Stbl3 chemically competent E. coli were transformed with In-Fusion products and plated on agar media with carbenicillin. Plasmids were extracted using the QIAprep Spin Miniprep Kit (Qiagen) and sequence-verified.

### Lentiviral transduction

To engineer cell lines with stably integrated constructs, L929s and C3H cells were transduced with lentiviral vectors in various combinations and sequences. To generate lentiviruses, Lenti-X HEK293T cells (Takara Bio) were plated in 6-well plates, and the next day, co-transfected the cells with the transfer plasmids plus two 2^nd^ generation packaging plasmids, pMD2.G and pCMVdR8.91.^114^ The Fugene 6 HD transfection reagent (Promega) or the TransIT-VirusGEN transfection reagent (Mirus Bio) were used to transfect the cells. Viral supernatants were collected 2 days after transfection and used immediately to transduce target fibroblasts. Most cell lines contained multiple integrated transgenes, and these were delivered either simultaneously by infection with multiple independently generated lentiviral vectors, or as a sequence of multiple rounds of transduction.

Target cells were first plated in 24-well plates (L929) or 6-well plates (C3H), and the next day, infected with freshly prepared lentiviral supernatants as described above. Typically, cells were exposed to a range of volumes of viral supernatant, from as low as 1 µL and up to 400 µL per well in the presence of 4 μg/mL polybrene (Sigma-Aldrich), to identify an optimal viral load. Most receiver cell lines were transduced such that about 1-2 integration events per cell occurred, as estimated based on the fraction of cells deemed positive for a given transgene. However, for sender cells, and for C3H cells, high expression levels were prioritized regardless of the number of integration events. Four to six days after infection, expression levels of transgenes were assessed by flow cytometry.

### Cell line selection

After at least 4 days following lentiviral infection, cells were lifted with TrypLE Express and transferred to a round-bottom 96-well plate. The cells were pelleted and resuspended in Ca^2+^-Mg^2+^-free PBS containing 2 mM EDTA and 2% FBS. When selecting for cell surface expression, cells were incubated with various fluorophore-conjugated antibodies, including an anti-myc AF647 (Cell Signaling Technology), anti-HA AF647 (Cell Signaling Technology), anti-Flag AF647 (R&D Systems), and anti-Her2 PE (BioLegend) antibody. Cells expressing the various transgenes were then isolated by cell sorting using a FACSAria II (BD Biosciences). To reduce some of the variability of the polyclonal cell populations, gates for sorting were set to only select cells in the middle portions of fluorescence distributions (when possible), thereby avoiding outliers on both ends.

For single cell isogenic clone selection in L929 cells, polyclonal receiver cell lines were first generated as described above. In a subsequent selection round, cells were again sorted by FACS and single cells were seeded into individual wells of a 96-well plate. Cells were allowed to grow for about 2 weeks and then screened for circuit performance compared to the average behavior of the parental polyclonal population. Single cell clones were then picked based on several criteria to be of use for further analyses, including baseline expression levels of the EGFP and mCherry targets and specific responsiveness to input combinations of synNotch ligands (CD19, nfGFP, and Her2).

### synNotch stimulation

Receiver cells with synNotch were stimulated with either magnetic beads coated with antibodies against Myc or HA epitope tags (Thermo Fisher), by co-culture with sender cells expressing ligands recognized by the synNotch extracellular domains, or by surface GFP ligand immobilized on a tissue culture plate. For experiments involving magnetic bead stimulation, receiver cells were plated in flat-bottom 96-well plates at a density of 16,000 cells per well. During the synNotch stimulation period, which ranged from as short as 0.5 days and up to 7 days, washed magnetic beads were added at density of 0.2 μL of bead slurry equivalent per well. Cells were passaged as normal throughout this period, and beads were re-added after each passage. To end synNotch stimulation, the gamma-secretase inhibitor, DAPT (TOCRIS), was added to the wells to a final concentration to 25 μM and the beads were removed using a magnetic plate. Note that DAPT was added to cultures in the control experiments where there was no synNotch stimulation.

For the experiment in which the relative timing of KRAB and D3L activity was varied, DAPT could not be used as it would inhibit both synNotch receptors. Instead, inhibition of the TetR-KRAB domain specifically was accomplished by addition of doxycycline to a final concentration of 1 μg/mL, while maintaining the anti-HA beads that stimulate synNotch-D3L. Similarly, inhibition of the CymR-D3L domain specifically was accomplished by addition of cumate (System Biosciences) to a final concentration of 60 μg/mL, while maintaining the anti-Myc beads that stimulate synNotch-KRAB. Cells were maintained in the presence of DAPT, doxycycline, or cumate, as appropriate, for at least 3 passages after removal of the beads.

For the experiment to test reactivation of the EGFP target gene with 5-Aza-2-deoxycytidine (5-Aza), cells were cultured in media containing 250 nM 5-Aza (Sigma-Aldrich), and cells were passaged daily until essentially all cells had died from 5-aza toxicity. Similarly, to test reactivation of the EGFP target gene via expression of TET1, cells were transduced with a lentiviral vector containing a TetR-TET1 (catalytic domain) construct following the initial establishment of memory. TetR-TET1 was constitutively expressed in the absence of synNotch stimulation or DAPT.

For the experiment to examine population bifurcation, cells were stimulated as above with magnetic beads for one day, allowed to recover for six days in the presence of DAPT, and then sorted into EGFP^high^ and EGFP^low^ groups using a FACSAria II (BD Biosciences). Three days after sorting, cells were restimulated with magnetic beads, in the absence of DAPT, to repeat the stimulation time course. For the experiment to test the delay time of mCherry activation in the inverter cascade, cells were grown during both stimulation and recovery periods in the presence of varying concentrations of Shield1 (Fisher Scientific), as indicated.

For L929 experiments involving co-culture with senders and receivers, cells were mixed and plated in round-bottom 96-well plates at a total cell density of 16,000 cells per well. Sender-receiver ratios were set to 4:1 at the beginning of the co-culture, or in the cases of multiple sender types simultaneously, 4:4:1 (sender1-sender2-receiver) or 3:3:3:1 (sender1-sender2-sender3-receiver). In the latter cases, total cell density was maintained at 16,000 cells per well. During the synNotch stimulation phase, cells were passaged every day for up to 4 days, or every 1-2 days for experiments with a 7-day stimulation period. This passaging regimen was developed empirically, and the goal was to maintain 50-100% confluency to increase the chances of cell-cell contact and synNotch stimulation. SynNotch stimulation was stopped by adding DAPT as above, and cells were maintained in the presence of DAPT for the remainder of the time course. However, for the experiment involving sequential stimulation with two different types of sender cells, the first stimulation was ended by FACS sorting to isolate the receivers. Following a recovery period, receivers were then co-cultured with the second set of senders as above. All experiments were performed in triplicate wells.

### Flow cytometry

At various timepoints during synNotch stimulation or recovery, cells were lifted with TrypLE and a portion were transferred to a round-bottom 96-well plate. Cells were then washed and resuspended in Ca^2+^- and Mg^2+^-free PBS containing 2 mM EDTA and 2% FBS. Fluorescence intensities of the resuspended cells were then assessed with either the BD LSR II or BD LSR Fortessa X20 with a high-throughput sampler. In addition to forward scatter and side scatter channels, fluorescence data was also collected across 4 channels, including BV 421 (TagBFP), FITC (EGFP), PE CF594 (mCherry), PE (mScarlet and PE-conjugated antibodies), and APC (iRFP670 and AF647-conjugated antibodies). Following acquisition, fluorescence data were first processed with FlowJo software (version 10, BD). Gates to determine the percent of cells positive or negative for EGFP or mCherry were established by untransduced (negative control) cell lines (Figures S1B and S5A).

### Sample prep for DNA methylation analysis

L929 cells were stimulated with magnetic beads as above, except cultures were scaled up in size to 6-well plate format. At various timepoints during synNotch stimulation and recovery, cells were lifted with TrypLE and about half the cells (∼5 x 10^5^) were transferred to tubes and flash frozen for subsequent processing in bulk. Genomic DNA was extracted using the Macherey-Nagel Nucleospin kit as described in the manufacturer’s protocol. Isolated gDNA was then digested with the NdeI enzyme (NEB) which maintains the KRAB-D3L target locus intact, including the SFFV promoter. Digestion was performed with 2-3 μg of total DNA and for 3 hrs at 37°C. Digested DNA was then cleaned with the QIAquick PCR Purification Kit (Qiagen) following the manufacturer’s protocols and a 50 μL elution volume. Digested DNA was then cytosine-converted using the Enzymatic Methyl-seq Conversion Module (NEB) following the manufacturer’s protocol. 100-200 ng of input DNA was used and formamide was used for denaturation, as suggested in the official protocol. The final converted DNA was purified with AmpureXP beads and processed directly for PCR amplification and long-read sequencing.

Following conversion and purification, two rounds of PCR were performed to amplify a 509 bp segment encompassing most of the SFFV promoter (containing 25 CpG sites) with unique barcodes introduced for de-multiplexing pooled libraries. In the first PCR round, NEB Q5U polymerase (NEB) was used according to the manufacturer’s recommendations and with the following parameters: this reaction was done in a 40 μL reaction volume using 10 μL of converted genomic DNA as input. Primers (designed with the Zymo Research Bisulfite Primer Design Tool) were diluted to a final concentration of 0.5 μM. The reverse primer contained custom unique index sequences for each sample. Thermal cycling was run as follows: 3 min at 98 °C, 35 cycles of 98°C for 8 sec, 62 °C for 30 sec, and 65 °C for 30 sec, and a final extension at 65 °C for 2 min. In the second PCR round, NEB Q5 polymerase (NEB) was used according to the manufacturer’s recommendations and with the following parameters: this reaction was done in a 40 μL reaction volume using 4 μL of undiluted unpurified amplicon from PCR 1 as input. Fresh primers (same as PCR round 1) were added to a final concentration of 0.5 μM. Thermal cycling was run as follows: 3 min at 98 °C, 25 cycles of 98°C for 8 sec, 59 °C for 30 sec, and 72 °C for 30 sec, and a final extension at 72 °C for 2 min. Amplified DNA was then run on a 1% agarose gel and bands ∼500 bp were extracted and purified using the Zymoclean™ Gel DNA Recovery Kit (Zymo Research) with an elution volume 10 μL. Extracted amplicons were then pooled (4 samples per pool) and analyzed by Nanopore long-read sequencing (Plasmidsaurus), which produced ∼5000 reads per pool.

### Chromatin immunoprecipitation with qPCR

L929 cells were stimulated with magnetic beads as above, except cultures were scaled up in size to T150 flasks. At various timepoints during synNotch stimulation and recovery, cells were lifted with TrypLE and 10 million cells were transferred and pelleted in 15 mL tubes. For cross-linking, pellets were washed once with ice-cold PBS supplemented with protease inhibitor cocktail (PI, Thermo Scientific), resuspended in 10 mL PBS+PI, and cross-linked by addition of methanol-free formaldehyde (Thermo Scientific) to a final concentration of 1%. Samples were incubated at room temperature for 10 min, after which cross-linking was quenched with 125 mM glycine for 5 min. Cells were pelleted, washed once in 10 mL PBS+PI, and again in 1 mL PBS+PI before transfer to 1.5 mL tubes and flash freezing.

For lysis, frozen pellets were thawed rapidly and resuspended in 1 mL cell lysis buffer (20 mM Tris-HCl pH 8, 85 mM KCl, 0.5% NP-40 and 1x PI in ddH_2_O) and rotated at 4 °C for 10 min. Nuclei were collected by centrifugation at 2,500 × g for 5 min at 4 °C, resuspended in 1 mL nuclear lysis buffer (1% SDS, 10 mM EDTA, 50 mM Tris-HCl pH 8 and 1x PI in ddH_2_O), and incubated on ice for 30 min to disrupt the nuclear membrane. Chromatin was sheared to an average size of ∼200 bp using a Covaris S2 sonicator in 1 mL Covaris tubes. For quality control, 5 µL of sonicated chromatin was reverse cross-linked overnight (RNase A, 30 min at 37 °C; proteinase K in 0.5 M NaCl, overnight at 65 °C), purified using a MinElute kit (Qiagen), quantified by Qubit (Thermo Fisher), and analyzed on a Bioanalyzer (Agilent). The remaining chromatin was snap-frozen until use. Prior to immunoprecipitation, lysates were clarified by centrifugation at maximum speed for 10 min at 4 °C.

For immunoprecipitation, clarified chromatin (40 µg of DNA) was pre-cleared with 15 µl of protein A+G magnetic beads (Sigma-Aldrich #16-663, pre-washed three times in nuclear lysis buffer) for 2 h at 4°C to remove non-specific binding. After applying to magnet, a 2% aliquot was reserved as input control. The remainder was diluted to 5 mL in ChIP dilution buffer (16.7 mM Tris-HCl ph8, 1.2 mM EDTA, 0.01% SDS, 1.1% Triton X-100, 167 mM NaCl and 1x PI in ddH_2_O) and incubated overnight at 4 °C in the presence of 8 µg of H3K9me3-specific antibody (Abcam #ab8898). The following day, 50 µL of protein A+G magnetic beads (pre-washed three times in ChIP dilution buffer) were added to each immunoprecipitation and incubated for 4 h at 4 °C. Beads were sequentially washed twice each with low-salt wash buffer (20 mM Tris-HCl pH 8, 150 mM NaCl, 2 mM EDTA, 0.1% SDS, 1% Triton X-100 in ddH_2_O), high-salt wash buffer (20 mM Tris-HCl pH 8, 500 mM NaCl, 2 mM EDTA, 0.1% SDS, 1% Triton X-100 in ddH_2_O), LiCl wash buffer (250 mM LiCl, 1% NP-40, 1% sodium deoxycholate, 1 mM EDTA, 10 mM Tris-HCl pH 8 in ddH_2_O), and TE buffer (10 mM Tris-HCl pH 8, 1 mM EDTA in ddH_2_O). Chromatin was eluted from beads in 50 µL freshly prepared elution buffer (1% SDS, 100 mM NaHCO_3_ in ddH_2_O) at 65 °C with shaking (1,000 rpm) for 30 min. Beads were magnetically separated, and the supernatant transferred to a new tube. A second elution was performed with an additional 50 µL elution buffer, and eluates were combined (final volume 100 µL). Cross-links were reversed by adding RNase A (0.5 µg/µL final) and incubating at 37 °C for 30 min with shaking, followed by addition of proteinase K (20 mg/mL) and NaCl (final 0.9 M) and incubation at 65 °C overnight for both ChIP and input samples.

DNA was subsequently purified using Qiagen MinElute PCR purification kit (Qiagen #28004). qPCR was performed in triplicate with locus-specific primers (promoter: F: CAGATGGTCACCGCAGTTT and R: ATAAGGCGCAGGGTCATTTC, gene body: F: GCTGACCATTAGCCCTGATTAT and R: CTTCAGCAGCTCCACATCAA) and primers to a negative control region (*GAPDH* locus, F: CTCTGGTAACTCCGCCTTTG, R: GATTACGGGATGGGTCTGAAC) in 1x SYBR green master mix (Roche Diagnostics, #50-720-3180) and run on a Real-Time Thermocycler (Bio-Rad CFX384) for 40 cycles. For fold enrichment analysis, triplicate Cq values were averaged for H3K9me3 ChIP and input samples separately. The following equation was used to normalize the Cq values of the ChIP DNA to the Cq value of the input DNA:

ΔCq_region of interest_ = Cq_ChIP-DNA_ – (Cq_Input-DNA_– log2[input dilution factor]), where input dilution factor = (fraction of input chromatin saved)-1 * input dilution factor pre-qPCR.

This calculation was repeated for the negative control (ΔCq_negative_). The final fold enrichment was calculated as 2^ΔΔCq^, where ΔΔCq = ΔCq_region of interest_ - ΔCq_negative_.

### Self-organizing spheroid assembly

Sender cells were first arrested to ensure they don’t outcompete receiver cells in the co-culture assays. To do so, sender cells in standard monolayer culture were incubated for 90 min at 37 °C in regular DMEM media containing 10 ug/mL mitomycin C (Abcam). In other experiments, sender cells, or control cells without ligands, were left untreated. Untreated and mitomycin-treated cells were then lifted with TrypLE (following washing of media) and resuspended in fresh media without mitomycin. Receiver cells, on the other hand, expressed a synCAM^107^ that disrupted adhesion to tissue culture plates, and as such they did not require enzymatic dissociation. Senders and receivers were then resuspended in DMEM with 4% FBS, which we empirically determined would enable receivers to self-assemble and grow as spheroids for up to 12 days in experiments without substantial overgrowth and cell death. Through serial dilution, we then added 30 untreated or 100 mitomycin-treated sender cells and 30 receiver cells into 96-well round-bottom ultra-low attachment plates (Corning #7007) in a final volume of 100 μL. Co-cultures were performed with at least 5 replicates. Co-cultures were then allowed to proceed unperturbed and undergo live-cell imaging daily by confocal microscopy.

We additionally prepared self-assembling spheroids for detailed time course assessment by light sheet microscopy in a similar manner with the following modifications. Mitomycin-treated senders and receiver cells were resuspended in Phenol Red-free Fluorobrite DMEM media (Thermo Scientific]) with 4% FBS and transferred to 96-well round bottom ULA plates as above (100 senders and 30 receivers). Importantly, receivers were a mixture of cells either labeled or not with an NLS-iRFP670 reporter. Labeled receivers were sparsely mixed in, making up 1/10 of the population (3 cells out of 30). This sparse receiver labeling enabled unambiguous segmentation and tracking within a densely packed, dynamically reorganizing 3D spheroid. This approach ensured that individual nuclear signals remained spatially resolvable throughout the imaging period, even as cells moved, divided, and reorganized within the aggregate. It also allowed more accurate assignment of fluorescence intensity traces to individual cells with lower risk of track-merging errors. Self-assembling spheroids were allowed to grow and self-organize unperturbed for 5 days after which they were carefully transferred by pipet to a sample holder for the Bruker TruLive 3D imager. This specific timepoint was selected to initiate light sheet imaging because it captured the period of greatest divergence in fluorescence dynamics between the KRAB-only (no memory) and KRAB+D3L (memory) conditions. After transfer, spheroids were manually inspected to ensure senders and receivers remained attached and intact. Sample holders were then overlaid with mineral oil (Thermo Scientific) and placed in the TruLive imager to undergo live-cell imaging for several days.

### Confocal microscopy of induced spheroids

Spheroids were directly imaged in the ULA cultures plates using the Opera Phenix automated spinning-disk confocal microscope (Revvity) with a 20× water-immersion objective lens (0.45 NA). Images were acquired daily for up to 9 days. Fluorescence data was acquired across 4 channels allowing determination of iRFP670 (some senders), TagBFP (some senders, and constitutively expressed in some receivers), EGFP (PCAD- receivers), and mCherry or mScarlet (PCAD+ receivers). Cross-sections were collected every 5-10 μM to create a total Z-stack 140 μM thick. Harmony software (Revvity) was used to operate the microscope and for image processing. Maximum intensity projections were generated with Harmony using the entire Z-stack at each time point. Harmony software was also used to apply contrast, brightness, and gamma values for digital image presentation (applied equally across samples). Additionally, raw image data was exported for further processing and quantification with the scikit-image and napari packages in Python.

### Light-sheet microscopy of induced spheroids

Time-course imaging of induced dual spheroids was performed using a Luxendo TruLive3D Imager (Bruker) equipped with a 25× water-immersion objective. This instrument is an inverted single-plane illumination microscopy platform that employs dual-sided light-sheet illumination. Two opposing excitation beams converge on the sample from opposite directions. The light-sheet was configured to a beam thickness of 1.9 µm and Z-stacks were acquired with a step size of 2 µm, spanning a total Z-axis range of 200 µm. The microscope also includes environmental control for temperature, humidity, and CO₂, enabling continuous imaging over multiple days. Samples were imaged every 10 minutes over a continuous 48-hour period. Samples were illuminated using 405 nm, 488 nm, 561 nm, and 645 nm laser lines to visualize TagBFP, EGFP, mScarlet, and iRFP670, respectively. To minimize spectral crosstalk, imaging was performed in two separate acquisition channels optimized for spectral separation. Fluorescence emission is collected through a high-numerical-aperture detection objective and then split between two cameras with sCMOS sensors. As the spheroids are not embedded in a matrix, the built-in sample tracking function of the LuxBundle software (Bruker) was enabled during acquisition to maintain the spheroids in focus and centered throughout the time course.

### Reprogramming fibroblasts toward a neuron-like state

To induce Brn2 + Ascl1 + Myt1L (BAM) expression in engineered C3H cells, we used two separate approaches. BAM factor activation through the synthetic memory circuit was achieved by seeding cells on a GFP-coated plate or by co-culturing cells with mitomycin C-treated L929 sender cells presenting cell-surface GFP ligand. GFP coated plates were prepared by first incubating standard 96-well tissue culture plates with Protein A (Sigma-Aldrich) at 20 µg/mL for 2 hours at room temperature, and then with a purified FC-GFP fusion protein at 5 µg /ml for 2 hours at room temperature. In both cases, the GFP ligand triggers anti-GFP synNotch receptors for both KRAB and D3L activation on the C3H cells. In a separate engineered cell line harboring Tet-ON induction, BAM factor expression was reversibly induced via doxycycline. In all cases, tissue culture plates were also coated with poly-D-lysine (Thermo Fisher) at 100 µg/mL for 30 min at 37 °C, followed by purified Laminin (Thermo Scientific) at 10 µg/mL for 30 min at 37 °C.

C3H cells were seeded at 10,000 cells per well in the laminin-coated (and some GFP-coated) 96-well plates in DMEM + GlutaMAX + 1 mM sodium pyruvate + 10% FBS. For the co-cultures, L929 senders were seeded at the same density for a 1:1 sender:receiver ratio (20,000 cells total). For the induction of BAM factor expression by doxycycline, cells were incubated with 2 ug/mL doxycycline. Cells were then grown in this medium for 5 days, with fresh media replacement every 2-3 days. At 5 days after initial plating, media was switched to N3 medium (DMEM/F-12 supplemented with Pen/Strep, N2, B27, and growth factors, hBDNF, hCNTF, hGDNF, hFGF2, all at 10 ng/mL)^115^ and cells were maintained in this medium for the remainder of the assay. Fresh media was added every 2-3 days. To stop synNotch input at 2 or 5 days, fresh media was added containing 25 µM DAPT (TOCRIS). Similarly, to stop synNotch input from L929 senders, cells were incubated in the presence of 25 µM DAPT and 1 µM AP20187 (Sigma-Aldrich), the latter of which induces apoptosis of the senders via iCaspase9. Doxycycline-based induction was stopped simply by washing cells and replacing with media lacking the molecule. Additionally, along with removal of doxycycline, cells were also incubated with DAPT as a control treatment.

### Immunofluorescence

At various timepoints after the initiation of BAM factor induction and recovery, media was removed and wells washed twice in PBS and fixed with 4% paraformaldehyde for 15 min at room temperature. Following fixation, cells were washed twice in PBS and permeabilized by three sequential incubations of 15 min with PBS containing 0.5% Triton X-100. After removal of the permeabilization buffer, samples were blocked in Blocking Solution (10 mg/mL BSA, 2 mg/mL evaporated nonfat milk, 0.5% Triton X-100, 1% DMSO) for 1 hr at room temperature. Primary antibody incubation was then performed for 1 hr at room temperature in Blocking Solution containing a 1:2000 dilution of the Tuj1 primary antibody against beta-tubulin III (GeneTex), or overnight at 4° C in Blocking Solution containing a 1:100 dilution of the primary antibody against VGLUT1 (GeneTex) or GAD65+GAD67 (Abcam). Samples were then washed three times for 10 min each in PBS containing 0.1% Tween-20. Secondary antibody incubation was performed for 1 hr at room temperature in Blocking Solution containing a 1:500 dilution of the goat anti-rabbit IgG Alexa Fluor 680 antibody (Thermo Fisher). Nuclear counterstaining was performed by adding DAPI during the secondary antibody incubation. Samples were subsequently washed three times for 10 min each in PBS containing 0.1% Tween-20. After the final wash, PBST was aspirated and replaced with 50 µL PBS for immediate fluorescence assessment on the Opera Phenix automated spinning-disk confocal microscope.

Fluorescence data was acquired on the Opera Phenix automated spinning-disk confocal microscope (Revvity) with 20× water or air objective lenses across four channels, with 405 nm, 488 nm, 561 nm, and 640 nm laser lines to visualize DAPI, GFP, and Tuj1 stained with the Alexa Fluor 680 antibody, respectively. Cross sections were collected every 4-6 μm to create total Z-stacks up to 20 μm thick. Harmony software (Revvity) was used to operate the microscope and for image processing. Maximum intensity projections were generated in Harmony using the entire Z-stack for each sample. Harmony software was also used to apply contrast, brightness, and gamma values for digital image presentation (applied equally across samples). For Tuj1-stained images, exported files were processed and analyzed using ImageJ software to segment DAPI-positive nuclei. Neuron-like cells were manually identified and counted based on the co-occurrence of: (1) the expression of the neuronal marker beta-tubulin III (Tuj1 stain), indicated by Alexa Fluor 680 fluorescence intensity above background; AND (2) the presence of thin Tuj1-positive neurite- or axon-like morphological protrusions. To calculate the percent of Tuj1+ neuron-like cells generated, the number of neuron-like cells was divided by the number of C3H cells initially plated.

### Quantification and statistical analysis

The numbers of replicates are indicated in the figure legends and method details. Flow cytometry raw data were first assessed using FlowJo (FlowJo LLC) to determine gates defining cells positive and negative for a given fluorescent marker. Data were then exported for further processing and visualization in Python, with the Jupyter Notebook and using Matplotlib and Seaborn packages. For line plots, confidence intervals are displayed as shaded areas under the lines (same color). For flow cytometry distributions, only one replicate is shown but they are representative of the set. One-dimensional distributions are plotted as kernel density estimate plots with Seaborn and with a bandwidth value (bw_adjust) of 0.5. Two-dimensional distributions are also plotted as kernel density estimate plots with Seaborn, but as a series of contour lines (15) of increasing darkness and with the lowest 5% of data omitted.

### DNA methylation analysis

FASTQ files were processed with a custom Python workflow taking advantage of the BioPython package.^116^ Sequence files were first de-multiplexed using the barcode sequences in a sense-aware manner, and resulting sequences were filtered for length greater than 509 bp (out of 549 bp expected including primers). For each filtered sequence, motifs surrounding each CpG site were used to scan the reads. The identity of all 25 CpG sites (in the reference sequence) were counted as being either cytosine or thymine. Any other nucleotide was discarded and assumed to be a mutation or PCR artifact. These site-specific C and T counts were used to generate single-read methylation heatmaps and for average methylation levels (average for each CpG across all reads and average across the entire promoter) that were then plotted using Seaborn.

### Induced spheroid volume calculation

For quantification of the size of induced PCAD+ spheroids in the self-organizing morphogenesis system, microscopy data from the Opera Phenix were exported and merged to form four-dimensional image stacks (time, x, y, z) for each fluorescence channel. Image analysis was performed in Python using scikit-image and SciPy.^117^ For each timepoint, the corresponding 3D subvolume of the mScarlet channel data was processed using a custom segmentation pipeline adapted from the scikit-image 3D watershed workflow. Briefly, raw intensity data were first smoothed with a Gaussian filter to reduce noise, and foreground–background separation was achieved by Otsu’s global thresholding approach, with an additive threshold offset to refine object boundaries. Local maxima in the distance map exceeding 70% of the global maximum were used as watershed seeds. The watershed transform was then applied to generate labeled segmentation volumes. For each timepoint, connected components were quantified using scikit-image. The largest connected object was assumed to correspond to the primary spheroid mass, and its volume was calculated as the product of voxel count and the physical voxel volume (10 x 1.196 x 1.196 μm^3^).

### Single cell tracking in induced spheroids

For single cell tracking of receiver cells within the induced PCAD+ spheroids, microscopy data from the TruLive Imager first underwent a pre-processing pipeline with various Python packages that included image transformation to correct for camera artifacts, such as mirroring between short and long cameras, conversion to Zarr format for efficient storage and parallel processing, and downsampling to reduce computational load, while preserving essential morphological features. Next, as the spheroids are not embedded in a matrix, they were free to tumble, jitter, and drift throughout the time course. To correct for this, inter-timepoint three-dimensional image registration was performed by calculating and aligning vectors between sender (TagBFP) and receiver (EGFP) mass centroids, ensuring accurate spatial correspondence across the time series. This was achieved with various Python packages, including Zarr and scikit-image.

Following image registration, individual cell segmentation and tracking based on the NLS-iRFP670 signal, were then performed using the Ultrack algorithm,^118^ which enabled automated identification of cells within the spheroid and temporal tracking of cell spatial trajectories throughout the three-day imaging period. Identified tracks were first filtered by duration, discarding tracks that were less than 60 hrs (out of 72). Note that many single cell tracks had to be discarded due to merging with another cell (preventing unambiguous tracking), or due to leaving the spheroid altogether at the edges. Both occurred stochastically. A few short/fragmented tracks were able to be merged using a linear assignment problem (LAP) based gap-closing step (Hungarian algorithm via scipy). Candidate links between track ends and starts were restricted to gaps of ≤2 frames and displacements of ≤15 voxels, the latter derived from the 95th percentile of observed frame-to-frame displacement and scaled modestly upward to accommodate the 2-frame gap allowance. Costs combined displacement and gap length, and the resulting global assignment was solved to avoid greedy mismatches. Merged tracks were visually validated in napari against segmentation masks prior to downstream analysis.

After identifying tracks of sufficient duration (and in a few cases merging short tracks), we focused on the final 48 hrs of the 72 hr time course. Segmented regions were used to sample the surrounding volume for EGFP and mScarlet channel intensity values. Masks were smoothed using a gaussian filter, then dilated. Images and movies were further processed and prepared for presentation using Python’s napari package. Displayed tracks in figures and movies were smoothed using a Savitzky-Golay filter fit to a second order polynomial to reduce noise while preserving features.

## SUPPLEMENTAL INFORMATION TITLES AND LEGENDS

**Figure S1, related to figure 2.**
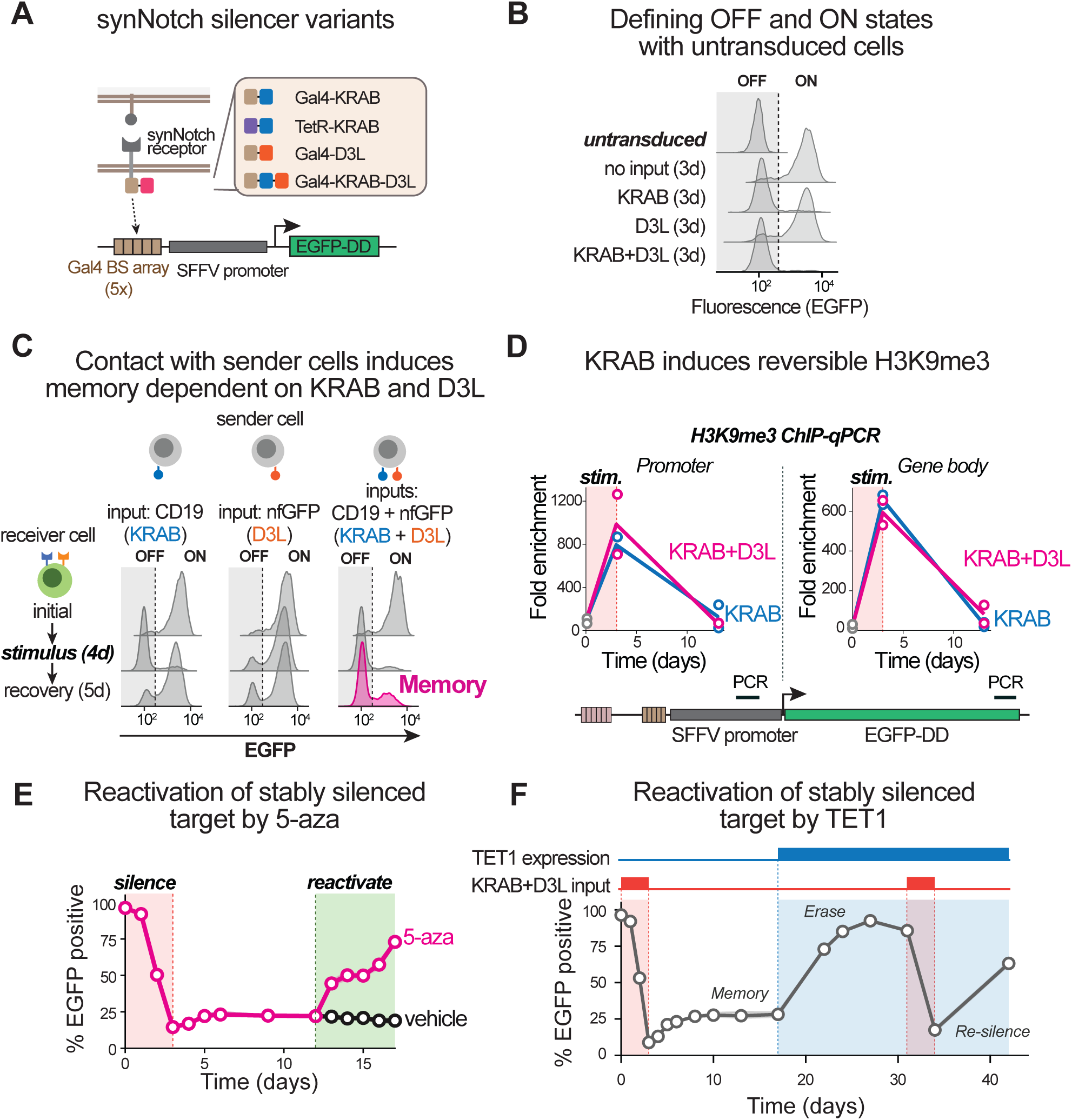
Inducible epigenetic memory through synNotch control of KRAB and Dnmt3L regulation, related to Figure 2. **(A)** Alternative configurations of the synNotch silencer receptors with different fusion combinations of DNA binding domain and chromatin regulators. **(B)** Flow cytometry distributions of untransduced L929 cells and engineered receiver cells after 3 days of synNotch stimulation. The untransduced cell background fluorescence was used to set the threshold for defining percent of cells expressing EGFP. **(C)** Flow cytometry distributions of receiver cells co-cultured with ligand-presenting sender cells as indicated. Cells were co-cultured for 4 days and allowed to recover for up to 7 days in the presence of DAPT. Distributions are representative of 3 technical replicates. **(D)** Time course of H3K9me3 levels at the engineered target locus in receiver cells that were transiently stimulated with anti-myc and anti-HA beads and recovered in the presence of DAPT. Individual data points (n = 2) are shown. The construct map below indicates the location of the PCR following ChIP. **(E)** Time course of the percent of cells expressing EGFP when they were initially incubated with anti-myc and anti-HA beads to stably silence EGFP, followed by treatment with 5-aza or vehicle. Data are from 3 technical replicates. Confidence intervals are too small to be visible. **(F)** Same as (B), but for presilenced receiver cells that were subsequently transduced with the human TET1 catalytic domain targeting the engineered locus, and then incubated a second time with anti-myc and anti-HA beads.

**Figure S2, related to figure 3.**
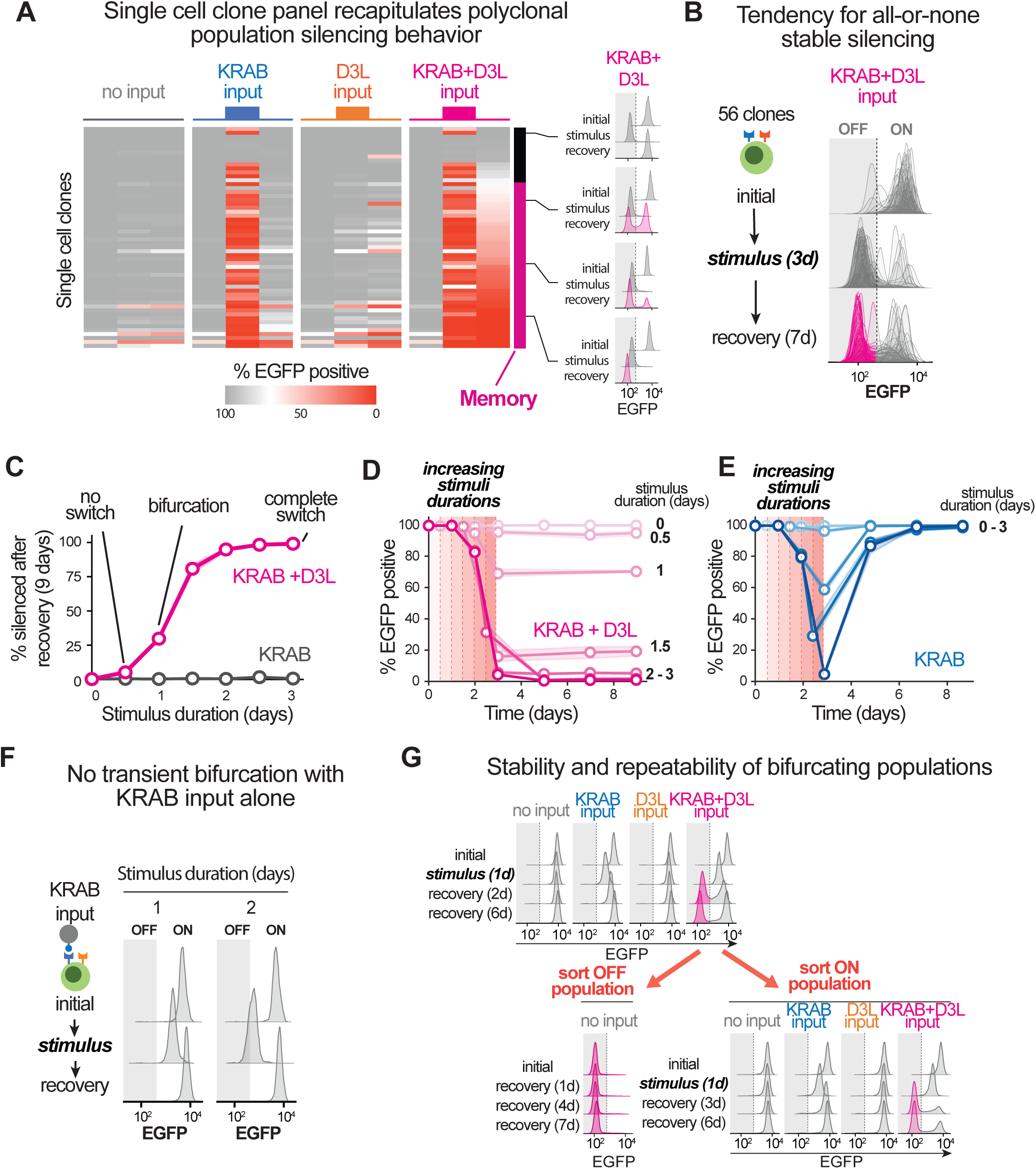
Bifurcation emerges upon KRAB and D3L stimulation in isogenic cell populations, related to Figure 3. **(A)** Heat map of the percent of cells expressing EGFP following the indicated stimulation types (colored bars above each group) for a panel of 56 randomly isolated single cell clones derived from the polyclonal population containing the circuit in Figure 2A. Each group of three columns corresponds to baseline, 3-day stimulation, and 7-day recovery periods. The black-pink to the right indicates clones that sustain the silenced EGFP during the recovery phase (Memory). Example fluorescence distributions from individual clones are shown to the right. Heatmap values are the average of 3 technical replicates. **(B)** Flow cytometry distributions of the panel of 56 monoclonal lines in (A). Distributions of all clones at each timepoint are overlayed. **(C)** The percent of cells in which EGFP was silenced following the recovery period (9 days) after a 3-day synNotch stimulation. Data shown are from 3 technical replicates of a single cell clone derived from the cell line in Figure 2. Confidence intervals are too small to be visible. **(D)** Time courses of the percent of cells expressing EGFP following the indicated stimulations. Receiver cells were incubated with beads to stimulate synNotch receptors and allowed to recover in the presence of DAPT. Data are from 3 technical replicates each. Confidence intervals are too small to be visible. **(E)** Same as (D). **(F)** Same as (B) but for a single receiver cell clone undergoing synNotch-KRAB stimulation for 1 or 2 days. **(G)** Same as (B), except that cells with high and low EGFP expression following KRAB+D3L stimulation were sorted by FACS at the end of a first recovery period. Cells were then restimulated with various inputs as indicated.

**Figure S3, related to figure 4.**
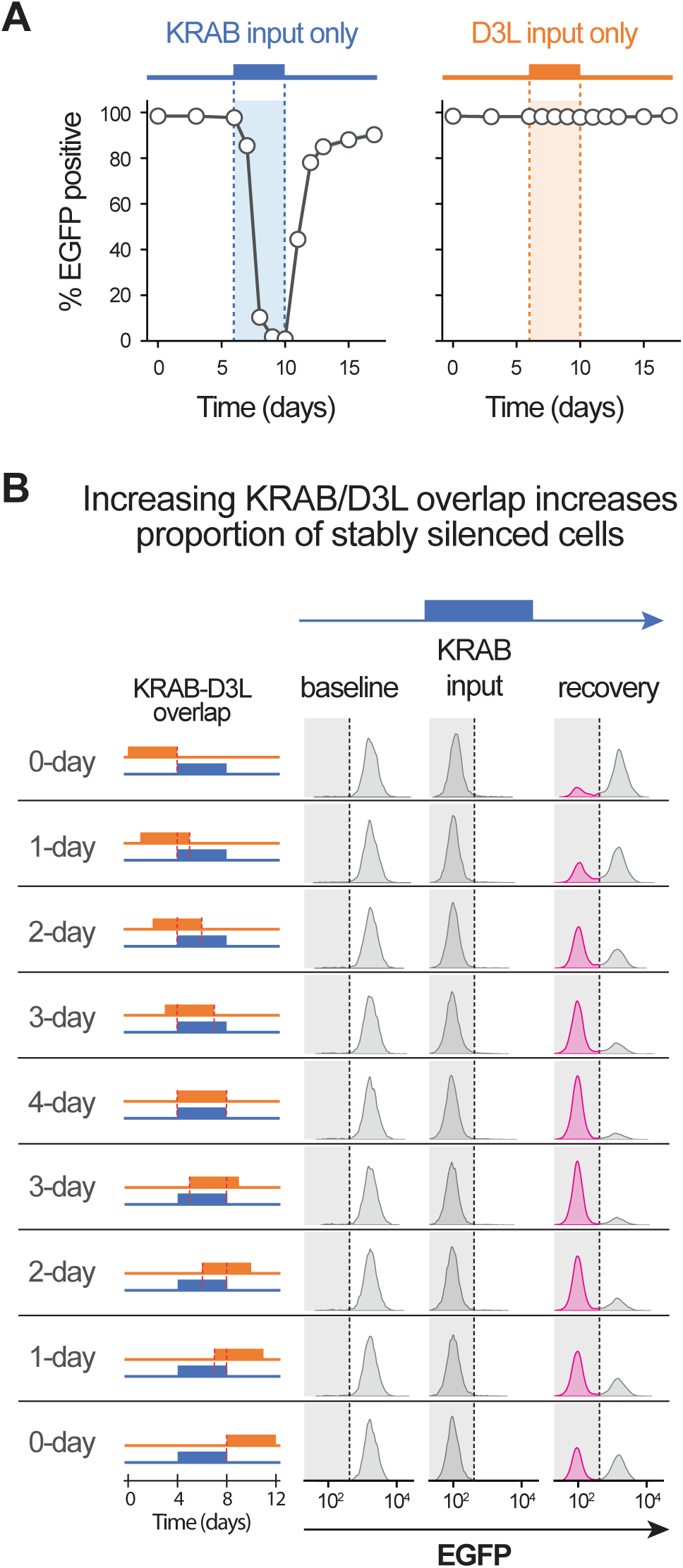
Strict requirement for KRAB and D3L co-stimulation to ensure cooperative stable state switching, related to Figure 4. **(A)** Time courses of the percent of cells expressing EGFP for either KRAB alone or D3L alone stimulation, as indicated by the colored bar and shaded area. Data for curves are from 3 technical replicates of a single cell clone. Confidence intervals are too small to be visible. **(B)** Flow cytometry distributions of receiver cells for varying stimulation regimens. Each column corresponds to a different timepoint, from prestimulation, to the end of the 4-day KRAB stimulation (blue bar above), to the end of the recovery period at 17 days. Each row indicates the KRAB-D3L overlap time along with a set of overlapping orange and blue bars illustrating the order of KRAB and D3L input. Distributions are representative of 3 technical replicates of a single cell clone.

**Figure S4, related to figure 5.**
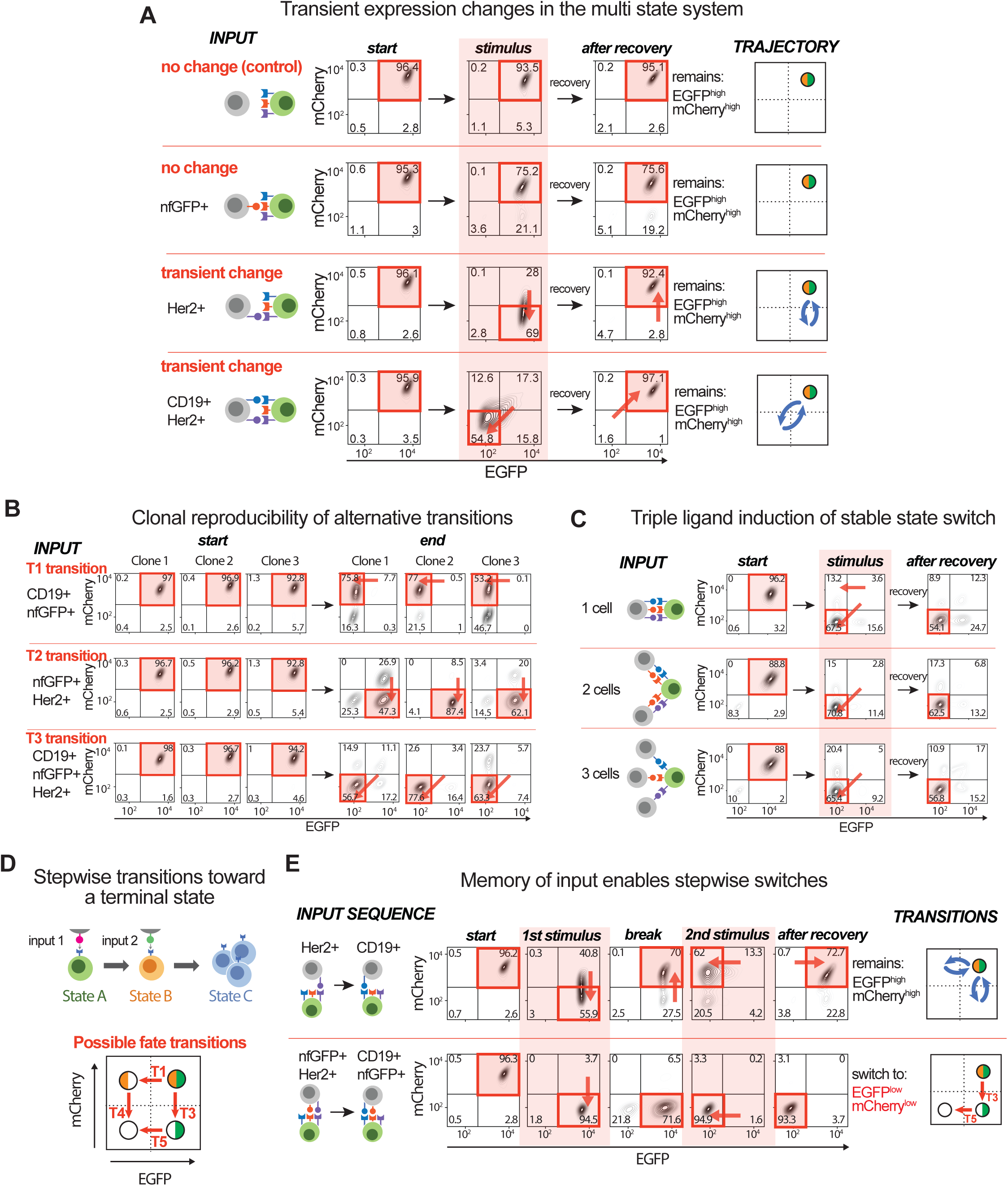
Transient, stable, and stepwise transitions between multiple alternative states are controlled by combinations of inputs, related to Figure 5. **(A)** Two-dimensional flow cytometry distributions of GFP and mCherry fluorescence. Receiver cells were co-cultured with different sender cells, as indicated in the ***INPUT*** column, for 7 days and then allowed to recover in the presence of DAPT for 7 days. Numbers are the percent of cells in each quadrant. Panels under the ***TRAJECTORY*** column summarize the dynamic changes in response to stimulation, all of which result in cells remaining in the baseline state. Distributions are representative of 3 technical replicates of a single cell clone. **(B)** Same as (A) but summarizing the start and end points for the three types of stable state transitions in three isogenic single cell clones harboring the circuit shown in Figure 5A. **(C)** Same as (A) but with alternative co-culture experiments for the T3 transition, with one, two or three sender cell types. **(D)** A schematic illustrating the possible stepwise transitions of the multipotent receiver cell toward a terminal cell state defined by EGFP and mCherry expression. Cells begin in the top right double-positive quadrant and can choose between two pathways based on the sequence of input combinations. T1 and T3 are similar to before, but T4 and T5 are introduced as new transitions that depend on the first transition. **(E)** Same as (A) but with sequential co-culture experiments for the T3 to T5 stepwise transition.

**Figure S5, related to figure 6.**
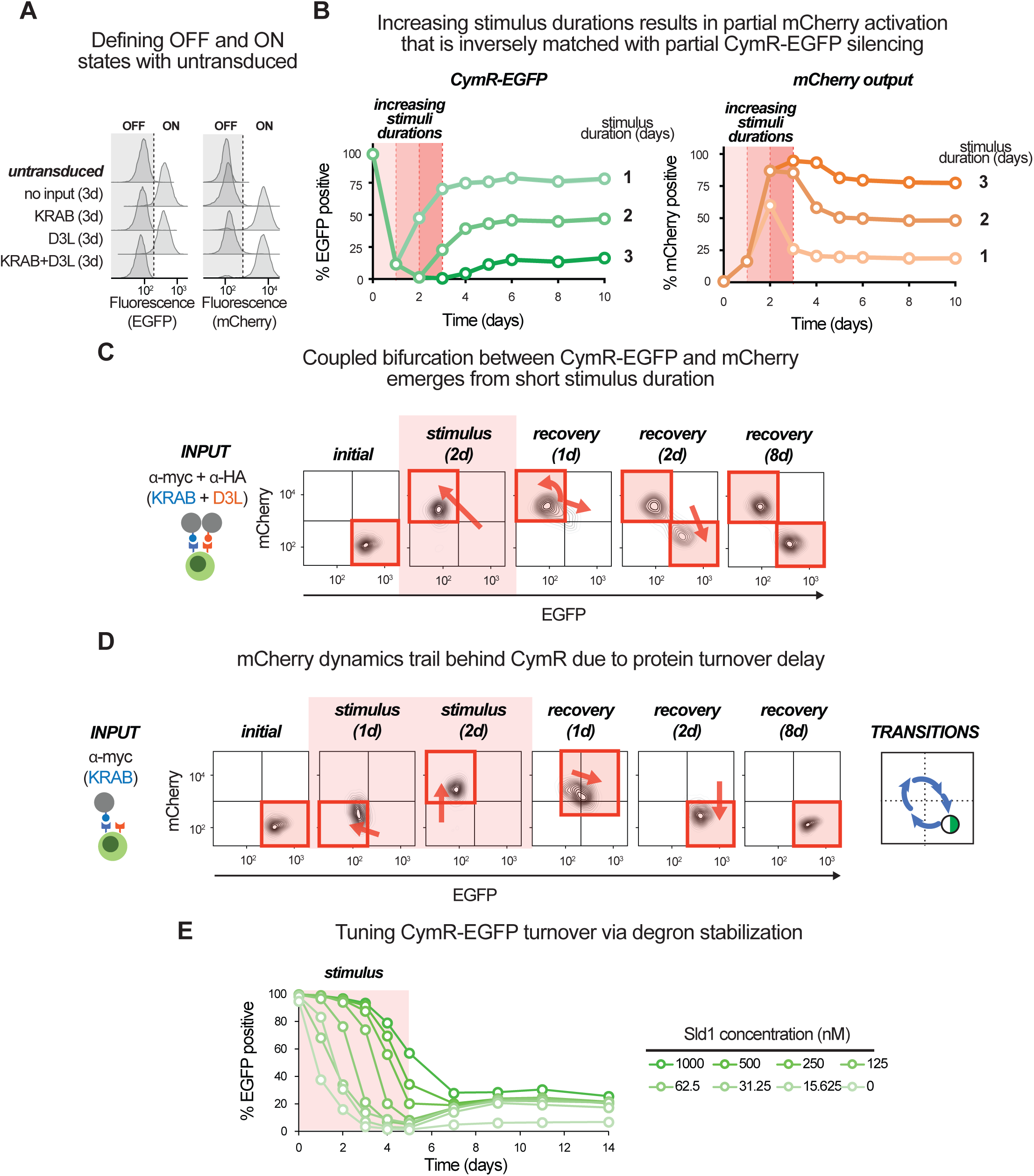
Activation is tightly coupled to silencing of CymR-EGFP and can also follow all-or-none bifurcation dynamics, related to Figure 6. **(A)** Flow cytometry distributions of untransduced L929 cells and engineered receiver cells after 3 days of synNotch stimulation. The untransduced cell background fluorescence was used to set the threshold for defining percent of cells expressing CymR-EGFP and mCherry. **(B)** Time courses showing the percent of cells expressing CymR-EGFP (left) or mCherry (right) in response to varying stimulus durations. Each red shaded area and dotted line indicates the increasing stimulus duration for each curve. Data for all curves are from 3 technical replicates of a receiver cell line derived from a single cell clone. Confidence intervals are too small to be visible. **(C)** Two-dimensional flow cytometry distributions of EGFP and mCherry fluorescence for receiver cells incubated with beads and allowed to recover in the presence of DAPT. Distributions are representative of 3 technical replicates. **(D)** Same as (C) but highlighting how mCherry lags behind CymR-EGFP during both silencing and reactivation phases in response to KRAB stimulation. **(E)** Time courses showing the percent of cells expressing CymR-EGFP following a transient stimulus and in the presence of varying amounts of the small molecule Sld1. Data for all curves are from 3 technical replicates. Confidence intervals are too small to be visible.

**Figure S6, related to figure 7.**
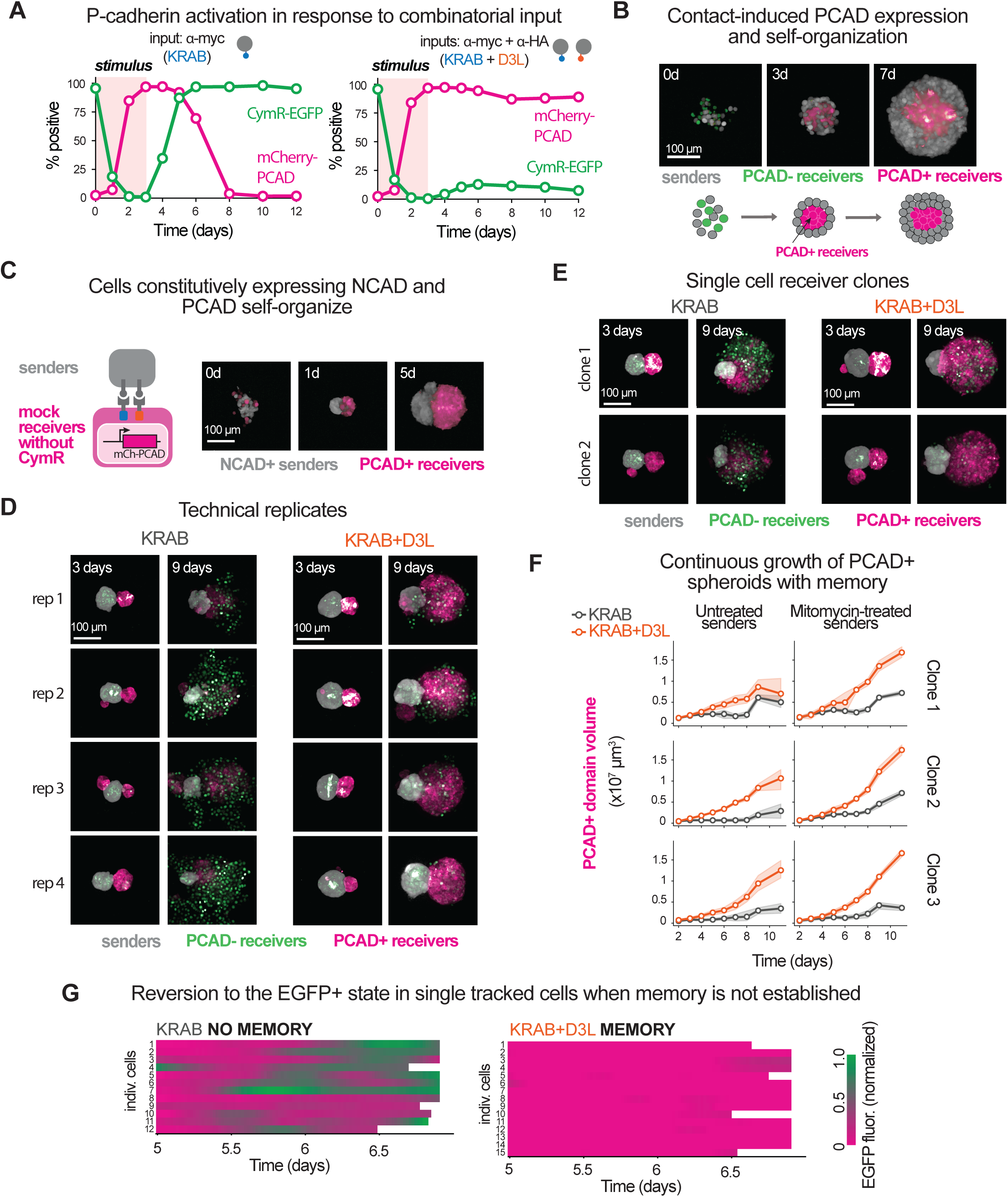
Stable activation of P-cadherin expression can be used to drive a stable morphogenic switch, related to Figure 7. **(A)** Time courses showing the percent of cells expressing CymR-EGFP or mCherry-P2A-PCAD in response to bead stimulation. Data for all curves are from 3 technical replicates of a receiver cell line derived from a single cell clone. Confidence intervals are too small to be visible. **(B)** Confocal microscopy images (maximum intensity projections) from an experiment where receiver cells were co-cultured with sender cells that do not ectopically express a cadherin in an ultra-low attachment well. The cartoon below summarizes the morphological dynamics. Images are representative of 4 technical replicates. **(C)** Same as (B) but with a co-culture experiment with NCAD+ senders and PCAD+ pseudo-receivers. Receiver cells harbor synNotch receptors but do not contain the CymR-EGFP construct that is the target of KRAB and D3L. Consequently, these cells constitutively express PCAD to mimic fully activated receivers. Images are representative of 4 technical replicates. **(D)** Same as (B) but with co-culture with NCAD+ sender cells that express CD19 (KRAB) or CD19+nfGFP (KRAB+D3L) ligands. Technical replicates are shown. **(E)** Same as (D). Different single cell clones are shown. **(F)** Time course of the size of the PCAD+ mScarlet+ domain during the self-organizing dual spheroid experiments. Volumes were measured from the confocal microscopy experiments shown in (D) and (E) plus an associated control experiment in which sender cells were not growth-arrested with mitomycin C (untreated). Shaded areas surrounding the curves are confidence intervals. **(G)** Heatmaps showing an extended set time courses of normalized EGFP fluorescence intensity for a panel of tracked cells within the induced spheroid structures (extended from Figure 7D). Note that some single cell tracks do not last the entire time course (white space) due to merging with another cell (preventing unambiguous tracking), or due to leaving the spheroid altogether at the edges.

**Figure S7, related to figure 8.**
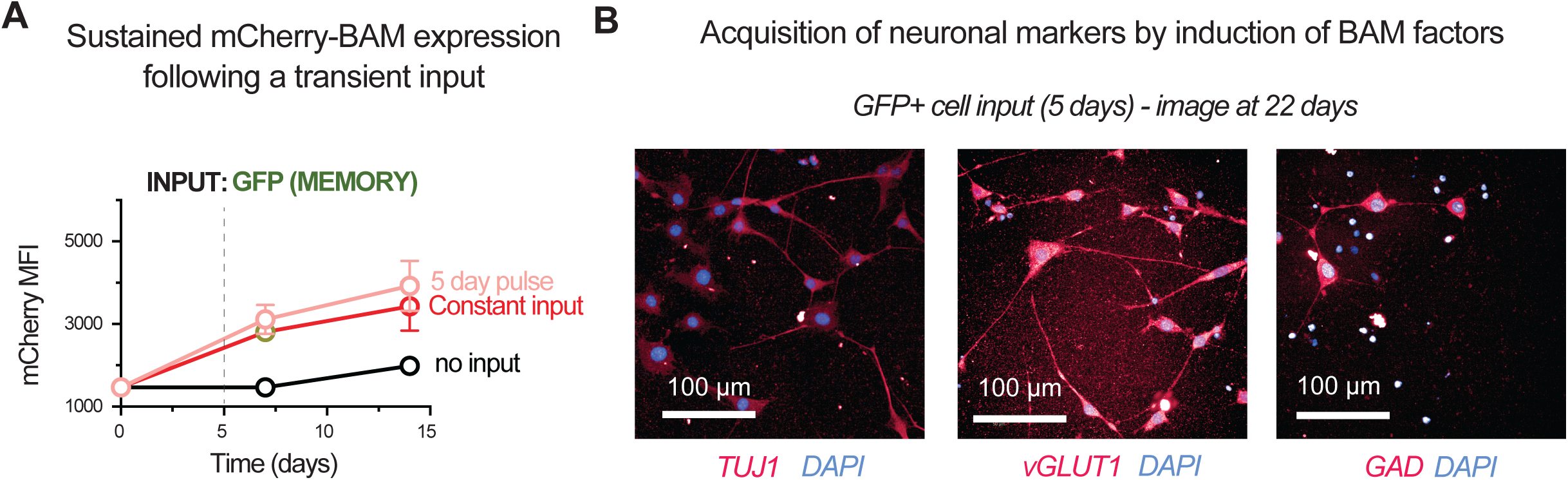
Epigenetic memory circuit enables a short input pulse to drive fibroblast reprogramming toward a neuron-like state, related to Figure 8. **(A)** Time courses showing the mCherry (reporter for BAM expression) mean fluorescence intensity (MFI) from flow cytometry analysis. Engineered C3H cells were stimulated with surface-immobilized GFP ligand for 5 days (input pulse) or 14 days (constant input). Data for curves are from 2 technical replicates. Error bars depict standard errors. **(B)** Confocal microscopy images showing cells stained with Tuj1, vGlut1, or Gad65+67 (GAD) antibodies at 22 days after the initiation of BAM factor expression via sender cell presenting GFP ligand.

**Video S1. Light sheet microscopy timelapse of individual tracked cells within induced spheroids, related to Figure 7E.**

Individual tracked cells are shown within spheroids induced with KRAB only (left) or KRAB+D3L (right) input. Expression of mScarlet-PCAD (magenta) and CymR-EGFP (green) is shown, and reversion events where cells return to the baseline EGFP+ state are labeled. The timelapse is from 5 to 7 days after the initiation of the co-cultures. The thin white lines that follow the tracked cells correspond to the trajectory from the prior 6 hrs. Note that the small number of green EGFP+ cells at the periphery of the MEMORY PCAD spheroid appear to have originated from receiver cells in the well that never came into contact with sender cells and thus never became EGFP-.

